# Hunger reconfigures a reward learning circuit into a memory competent mode

**DOI:** 10.64898/2026.09.17.752172

**Authors:** Annie Park, Bhagyashree Senapati, Nino Mancini, Youchong Zhang, Juliana Choi, Ian Cone, Christoph D. Treiber, Rui Ponte Costa, Lisa Fenk, Salil Bidaye, Scott Waddell

**Affiliations:** Centre for Neural Circuits and Behaviour, University of Oxford, Oxford, OX1 3SR, UK; Max Planck Florida Institute for Neuroscience, Jupiter, FL, USA; Active Sensing, Max Planck Institute for Biological Intelligence, Martinsried, Germany

## Abstract

Internal states such as hunger dynamically reshape activity across circuits to support resource seeking. Neuromodulation provides a means of controlling such physiological properties of neurons, but how this flexibility regulates memory networks remains unclear. Here, we describe how hunger reconfigures a dopaminergic food-reward circuit between two physiological modes that support memory formation, in *Drosophila*. Starvation suppresses baseline dopaminergic activity through peptidergic signalling, enabling reward-evoked, large-amplitude dopamine neuron spikes to reinforce learning. This spiking mode can be engaged by sugar consumption and persists beyond feeding, reflecting the fly’s satiety state.Persistent dopaminergic large-amplitude spiking reinforces learning and transitions the memory network into a mode that prioritizes consolidation of recently acquired memories. The transition from hunger to satiation is also reflected in the activity of postsynaptic output neurons that shift from a tonic, decorrelated mode that is responsive to dopamine into a bursting, correlated mode in which further dopaminergic modulation is occluded. Therefore, nutrient deprivation reconfigures the memory circuit into a learning competent dopamine-receptive mode, which persistent reinforcing dopamine then switches into a satiated mode driving memory consolidation. Together these processes mechanistically intertwine state-dependent reward-signalling with subsequent memory stabilization through transitions in physiological mode.

## Main

Hunger changes physiology throughout the brain, driving neural circuits to adaptively promote resource seeking and learning. In the compact *Drosophila* nervous system, neuromodulation provides a powerful means of achieving such state-dependent flexibility, expanding the functional repertoire of circuits and enabling complex behavior with relatively few neurons^1^. Although elegant studies in a handful of model systems have uncovered detailed molecular and physiological mechanisms by which neuromodulators reconfigure circuit function ^2–4^, it remains unclear how internal state adaptively modifies circuit function to support state-dependent learning.

It is now possible to address this gap with unprecedented mechanistic detail in *Drosophila*, owing to the completion of connectomes and single-cell transcriptomes. However, despite these resources it remains a central challenge to determine how adaptive physiology emerges from cellular and molecular components, and how it is altered by internal state to support behavior. Here we investigate the nature of a physiological transition that occurs during sugar learning, a shift from hunger to satiety as flies consume a sucrose reward. We first determined physiological and molecular mechanisms driving this transition, then tested how these mechanisms contribute to memory formation. Surprisingly, we identified an unprecedented degree of physiological flexibility during this transition, which permits state-dependent learning and subsequent memory consolidation.

### Food reward dopaminergic neurons are modulated by hunger state

Distinct dopaminergic neuron types encode different rewards. Although their anatomical and molecular heterogeneity suggests each type could engage distinct physiological computations^5–8^, this has largely been unexplored. The α1 protocerebral anterior medial dopaminergic neuron type (α1 DANs), which consists of only eight neurons per hemisphere, is required for long-term sugar-reinforced memory (Fig. 1a; Supplementary Fig. 1a) ^9,10^. Formation of long-term sugar-reinforced memories depends on the nutrient value of the reward, and requires that the flies are hungry prior to training ^9,11^, indicating that the transition from hunger to satiety may be a critical regulator of α1 DAN function.

**Figure 1:**
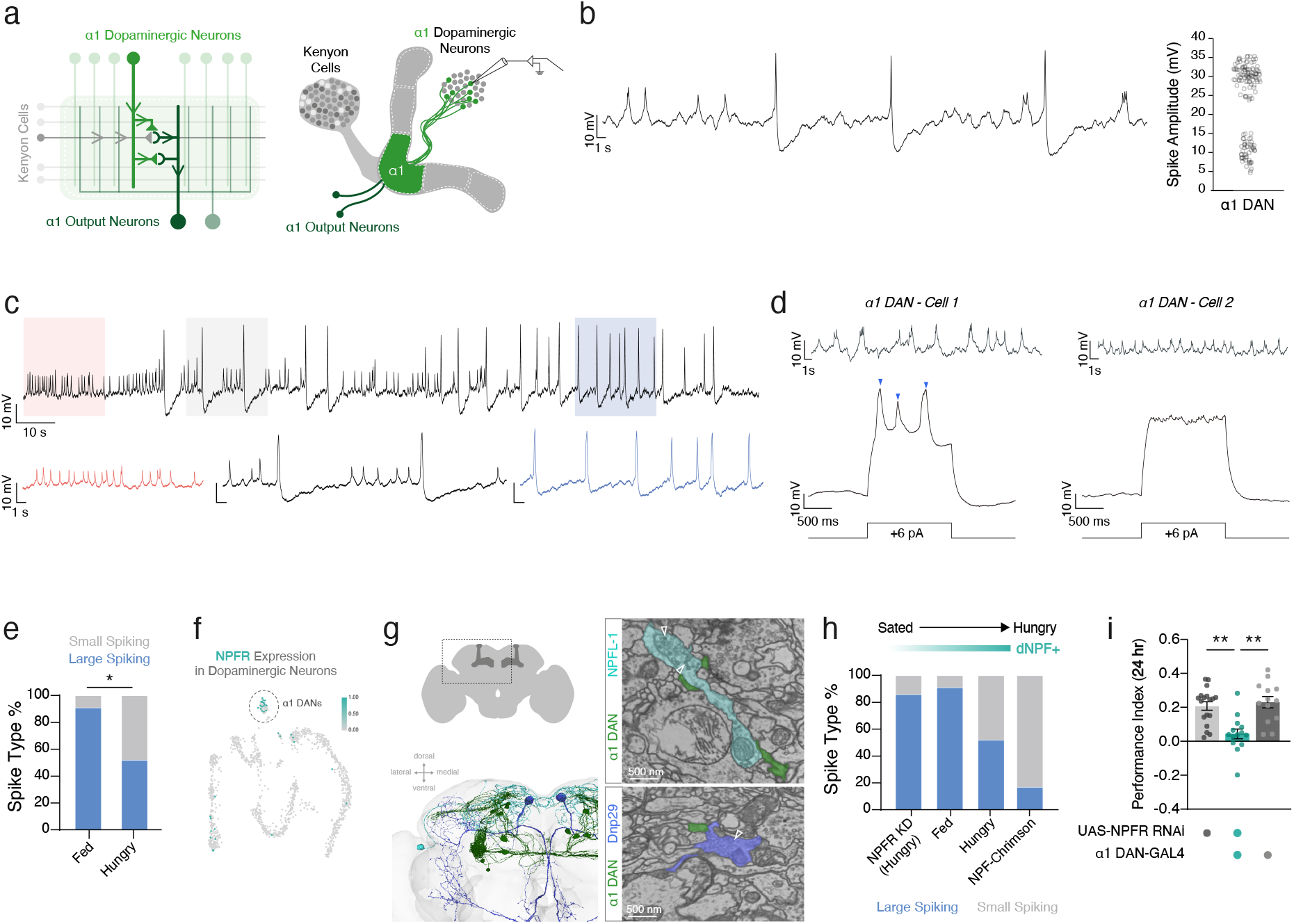
NPF couples hunger state to spiking properties in a1 DANs. **a,** Schematic of the Mushroom Body showing α1 dopaminergic neurons (α1 DANs), and α1 Mushroom Body Output Neurons (α1 MBONs). α1 DANs synapse onto α1 MBONs and form polyadic synapses with αβ Kenyon cells and α1 MBONs. **b,** Representative spontaneous activity trace of a single α1 DAN exhibiting small and large spike amplitudes. Quantification of spike amplitudes from a single neuron. **c,** Representative spontaneous activity trace of a single α1 DAN transitioning from a small to large amplitude spiking mode. **d,** Left, example small-amplitude spiking α1 DAN that generates large-amplitude spikes (blue arrowheads) with depolarizing current injection. Right, example α1 DAN unable to generate large-amplitude spikes. **e,** Proportion of α1 DANs capable of producing large-amplitude (blue) vs. small-amplitude spiking only (grey) (Fisher’s exact test; p=0.031, n=11,25 neurons – one per fly). **f,** UMAP visualization of dopaminergic neurons by relative Neuropeptide-F receptor (*NPFR*) expression (cyan). **g,** Left, schematic of the imaged window relative to the whole brain (top) and EM reconstruction of α1 DANs (dark green) and NPF+ neurons (NPFL-1, cyan and Dnp29, blue) (FlyWire, FAFB). Right, representative EM images of contact sites between NPF+ neurons and α1 DANs; arrowheads mark dense-core vesicles. **h,** Proportion of α1 DANs able to produce large-amplitude spikes increases with satiation; increased relative NPF reduces this likelihood (Fisher’s exact test; p=0.0009) (n=7,11,25,12 neurons – 1 neuron per fly, left to right). **i,** Knockdown of *NPFR* in α1 DANs impairs 24h memory performance (One-way ANOVA with Tukey’s multiple comparisons, n=18,14,12 groups of flies, p=0.0003 & p<0.0001).

To investigate how hunger state modulates the capacity of α1 DANs to instruct long-term memory formation, we performed in-vivo whole-cell patch-clamp recordings of α1 DANs from flies fed ad-libitum. We took this exploratory approach to preserve natural individual variation of hunger, permitting examination of the underlying variation in physiology^12^. α1 DANs exhibited different spiking patterns sometimes switching between patterns, with individual cells exhibiting the capacity to produce two spike amplitudes at the same time, small (∼10 mV) and large (∼30 mV) (Fig. 1b,c). Neurons spontaneously firing small-amplitude spikes could be made to fire large-amplitude spikes with depolarizing current injection (Fig. 1d, left). Current injection did not elicit large-amplitude spikes in some α1 DANs, indicating that formation of large-amplitude spikes is tightly regulated (Fig. 1d, right).

Although we considered that different spike patterns could arise from cell-type-specific properties of the three anatomical α1 DAN subtypes ^13^, observing spontaneous switching between spiking patterns within a single neuron argued against cell-intrinsic specialization. We therefore tested whether α1 DAN spike patterns could be dynamically regulated by hunger state-dependent neuromodulation, as α1 DANs are critical for forming sugar rewarded memories. To assess the ability of the α1 DANs to produce large-amplitude spikes, we injected sweeps of depolarizing current and quantified whether large-amplitude spikes emerged. Hungry flies fed sucrose immediately before recording showed an increased probability that α1 DANs could generate large-amplitude spikes, compared to starved flies (Fig. 1e). Importantly, spike frequency did not change between hunger states (Supplementary Fig. 1b), indicating that hunger modulates α1 DAN spike pattern rather than their general level of activity.

To identify the neuromodulatory basis of spike pattern emergence, we used single-cell transcriptome data to identify neuropeptide receptors enriched in α1 dopaminergic neurons. These analyses revealed expression of *Drosophila* neuropeptide F receptor (NPFR) in α1 dopaminergic neurons (Fig. 1f). *Drosophila* Neuropeptide F (dNPF) is the *Drosophila* homologue of mammalian neuropeptide Y and has been implicated in hunger-state signalling, motivational control of feeding and memory expression ^14,15^. Complementary analysis using the whole-brain connectome revealed areas of contact between dNPF-expressing neurons and α1 dopaminergic neurons, with dense-core vesicles in dNPF neurons positioned near these contact sites (Fig. 1g; Supplementary Fig. 1c). We therefore tested whether hunger state as represented by dNPF signalling regulates the capacity of α1 dopaminergic neurons to generate large-amplitude spikes. We first imposed an artificial hunger state using optogenetic stimulation of dNPF-producing neurons in fed animals ^14^, while recording α1 dopaminergic neurons. This manipulation reduced large-amplitude spike generation, as measured by depolarizing current injection, emulating the effects of starvation (Fig. 1h). To specifically implicate dNPF signalling within α1 dopaminergic neurons, we knocked down *NPFR* selectively in α1 dopaminergic neurons using RNAi. Strikingly, *NPFR* knockdown increased large-amplitude spike generation, mimicking the satiated state.

Together, these results demonstrate that dNPF signalling couples hunger state to spike pattern generation in α1 dopaminergic neurons, possibly by setting the dynamic range for reward-evoked activity. Reduced dNPF signalling in the satiated state increases spontaneous large-amplitude spikes, reducing the relative difference between baseline and reward-evoked dopaminergic neuron activity. In line with this idea, *NPFR* knockdown in α1 dopaminergic neurons impaired sucrose-reinforced long-term memory (Fig. 1i; Supplementary Fig. 1d), indicating that hunger-dependent dNPF signalling establishes a physiological mode permissive for reward-evoked long-term memory formation.

### Large-amplitude spikes provide an instructive signal for memory formation

Having established that hunger regulates the physiological mode of α1 dopaminergic neurons, we wondered whether spike modes also differ in their capacity to instruct memory formation. We therefore identified a circuit that could potentially allow precise control of different spike patterns in a physiologically relevant manner. Connectome analysis revealed a single cholinergic inter-neuron (PRW.SMP.3 or PhG1a & PhG1c) that receives direct cholinergic inputs from sweet-taste gustatory receptor neurons (GRNs) (e.g. Gr43a-expressing GRNs) and projects to the dendrites of the α1 dopaminergic neurons (Fig. 2a; Supplementary Fig. 2a)^16,17^. We next varied the frequency of optogenetic stimulation of sweet-taste GRNs to elicit small or large-amplitude spikes in α1 DANs (Fig. 2b,c; Supplementary Fig. 2b). These frequencies were taken from a previous observation that long-term memory formation depends on GRN stimulation frequency^18^. We confirmed this by observing that robust short- and long-term memory formation was restricted to stimulation frequencies that elicited large-amplitude α1 DAN spikes, despite equivalent total light delivery across conditions (Fig. 2d).

**Figure 2:**
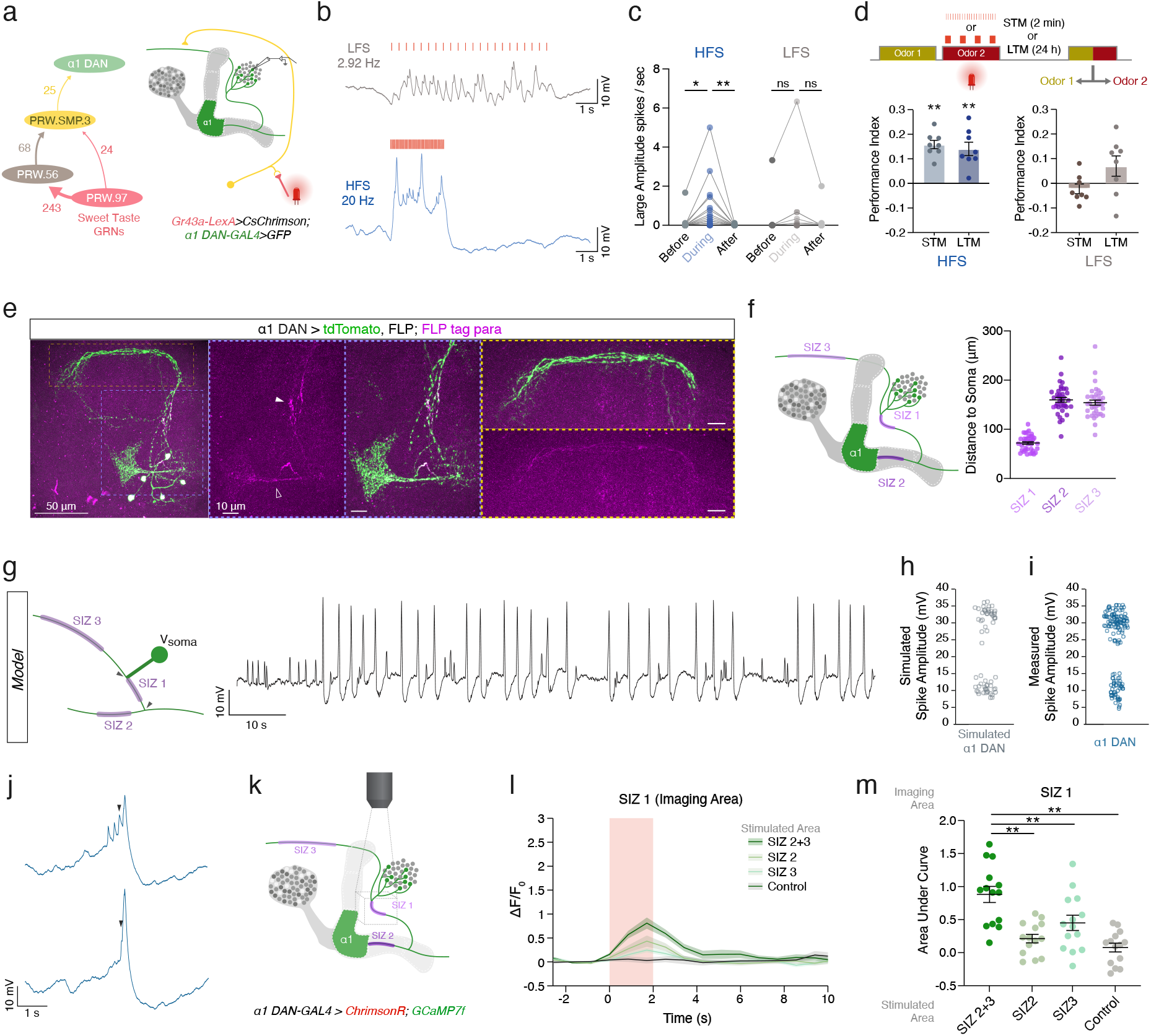
Multiple spike initiation zones enable distinct activity modes in a1 dopaminergic neurons. **a,** Left, Sweet taste inputs (red, PRW.97) and secondary projections (tan, PRW.56; yellow, PRW.SMP.3), onto α1 DANs with synapse numbers (FlyWire FAFB). Right, GRNs (*Gr43a-LexA*) were light-activated during α1 DAN patch-clamp recordings. **b,** Low-frequency stimulation (LFS; 2.92 Hz, top) generated small spikes; high-frequency stimulation (HFS; 20 Hz, bottom) elicited large-amplitude spikes. **c,** HFS evoked large-amplitude spikes (n=20 neurons – 1 neuron per fly with repeated measures, Friedman test with Dunn’s multiple comparisons, Before vs. During p=0.0126, During vs. After p=0.0049) but not LFS (n=20 neurons – 1 neuron per fly with repeated measures, Friedman test with Dunn’s multiple comparisons, Before vs. During p=0.7052, During vs. After p=0.8816). **d,** Flies learned odor pairings with HFS (n=8 groups of flies, one-sample t-test STM p<0.0001, LTM p=0.0013), but not LFS (n=8 groups of flies, STM p=0.2868, LTM p=0.1342). **e,** Endogenous para (magenta) and membrane-bound tdTomato (green) in α1 DANs. **f,** Distance from each spike-initiation zone (SIZ) to the soma (Friedman test with multiple comparisons correction; SIZ 1 vs. SIZ 2 p<0.0001, SIZ 1 vs. SIZ 3 p<0.0001, SIZ 2 vs. SIZ 3 p>0.999; n=37 brains, mean ± s.e.m.). **g,** α1 DAN multi-compartment biophysical model; recording site from V_soma_, with representative voltage trace (right). **h,** Modelled spike-amplitude distribution, recapitulating **i,** experimentally observed distribution. **j,** Traces of small spikes (black arrowhead) ramping up to a large-amplitude spike. **k,** Holographic optogenetic stimulation of SIZ 2/SIZ 3 during calcium activity recording at SIZ1 (*α1 DAN-GAL4>ChrimsonR; GCaMP7f*). Distal SIZ2 and SIZ3 were stimulated either individually or simultaneously. **l,** Mean calcium responses (ΔF/Fo) at SIZ1. Mean + s.e.m.. **m,** Area under the ΔF/Fo curve (n=14 brains; repeated-measures one-way ANOVA with Geisser-Greenhouse correction, F(1.948,25.32) = 19.28, p<0.0001, ε=0.6493, partial η²=0.597; Šídák’s multiple comparisons test, SIZ 2+3 vs. SIZ2 p=0.0014, SIZ2+3 vs. SIZ3 p=0.0051, SIZ2+3 vs. Control p=0.0002).

### Coincident activation of multiple SIZ underlies large spike amplitude formation

Given the importance of large-amplitude spikes for memory formation, we investigated the mechanism underlying spike amplitude generation in α1 dopaminergic neurons. We began by testing whether all spike amplitudes generated were sodium spikes by bath perfusing tetrodotoxin (TTX) while patch-clamp recording from α1 dopaminergic neurons. Addition of TTX suppressed all spike amplitudes even with depolarizing current injection (Supplementary Fig. 2e,f). Given that all spike amplitudes required voltage-gated sodium channels, we next examined if individual α1 dopaminergic neurons might contain multiple spike initiation zones (SIZ). We hypothesized that differential spike amplitude at the soma could be determined by the relative position of the initiating SIZ, e.g. large-amplitude spikes are initiated at a proximal SIZ because they undergo less attenuation. Using a FlpTag approach, we visualized GFP-tagged endogenous voltage-gated sodium channels in α1 dopaminergic neurons, which are encoded by *paralytic* (*para*) in *Drosophila*^19^(Fig. 2e; Supplementary Fig. 2c,d). Across brains, we observed three distinct regions of enriched para expression in α1 dopaminergic neurons, indicating three SIZs, which had different distances from the soma (Fig. 2f).

While these findings implicated SIZ position relative to the soma as a determinant of spike amplitude, it was unclear why spikes initiated at the distal SIZ 2 and SIZ 3 did not consistently recruit the intervening proximal SIZ 1. Moreover, connectomic analysis revealed relatively fewer synaptic inputs near SIZ 1 (Supplementary Fig. 2g,h), raising the question of how the proximal SIZ could be activated. To address these questions, we first constructed a biophysically detailed, multicompartment Hodgkin–Huxley model containing three SIZs positioned according to their morphologically identified locations (Fig. 2g). The model recapitulated the multiple spike-amplitude modes observed in our recordings (Fig. 2h,i).

Notably, patch-clamp recordings frequently revealed small-amplitude spontaneous spikes immediately preceding a large-amplitude spike (Fig. 2j), suggesting that activity initiated at the distal SIZs could recruit the proximal SIZ, similar to a mechanism previously described in locust neurons^20^. Consistent with this interpretation, our model generated small-amplitude spikes that were initiated independently at either distal SIZ (SIZ 2 or 3), which attenuated electrotonically as they propagated towards the soma. By contrast, near-coincident activation of SIZ 2 and SIZ 3 could recruit activation of SIZ 1, resulting in a large-amplitude somatic spike (Supplementary Fig. 2i). Together, these results support a conceptual framework in which interacting SIZs produce a physiologically distinct spiking mode that is relevant for learning.

To test our proposed mechanism of coincident SIZ integration, we used optogenetic holographic stimulation to precisely activate SIZs in α1 dopaminergic neurons, while recording calcium activity across the neurites of the α1 dopaminergic neurons (Fig. 2k). Although we implicated sodium spikes as the main mechanism underlying different spike amplitudes, we used calcium as a readout for local activity along the neurites. Simultaneous optogenetic stimulation of SIZ 2 and SIZ 3 while imaging from SIZ 1 resulted in a greater calcium response than optogenetically stimulating SIZ 2 or 3 alone (Fig. 2l,m), supporting our model that coincident activation of SIZ 2 and 3 facilitates proximal SIZ activation.

Taken together, these data support a mechanism underlying a specialized spiking mode, which is required for learning. Hunger state gates this process, as hungry flies have a diminished ability to produce large-amplitude spikes likely due to neuromodulatory control of SIZ 1 (e.g. reduced surface-level *para* expression). We propose that as flies begin to consume sugar, α1 dopaminergic neurons rapidly integrate changes in reward and nutrient state ^21^, increasing the excitability of SIZ 1, enabling it to generate large-amplitude spikes. Although we have not yet resolved whether somatic spike amplitude determines magnitude of dopamine release, the selective association between large-spike-generating stimulation regimes and memory formation indicates that the ability to engage this spike mode is essential for learning.

### Hunger state modulates α1 MBON bursting mode by enhancing electrical coupling

Having established that hunger state regulates α1 dopaminergic neuron spike mode engagement, we next investigated whether α1 dopaminergic neuron spiking modes differentially influence downstream circuit function. Each compartment in the Mushroom Body contains dendritic terminals from one to eight output neurons (MBONs), which project axonal terminals both inside and outside of the Mushroom Body to bias activity towards approach or avoidance ^13,22–25^. While previous work has largely focused on hunger-state-dependent changes in odor-evoked responses ^23,26–29^, it is unclear if internal state modulates intrinsic physiological properties of MBONs to facilitate memory formation.

In our patch recordings of α1 MBONs, we observed that spiking patterns also changed depending on the animal’s hunger state. Starved animals exhibited tonically spiking α1 MBONs, whereas those in fed animals were regularly bursting (Fig. 3a,b; Supplementary Fig. 3a-d). Because spiking mode reshapes network-level properties such as the degree of coincident spiking ^30,31^, we asked whether coincident spiking between ipsilateral α1 MBONs was hunger regulated. Dual whole-cell patch-clamp recordings of ipsilateral α1 MBONs exhibited greater spike synchronization in fed flies than starved (Fig. 3c-e; Supplementary Fig. 3e).

**Figure 3:**
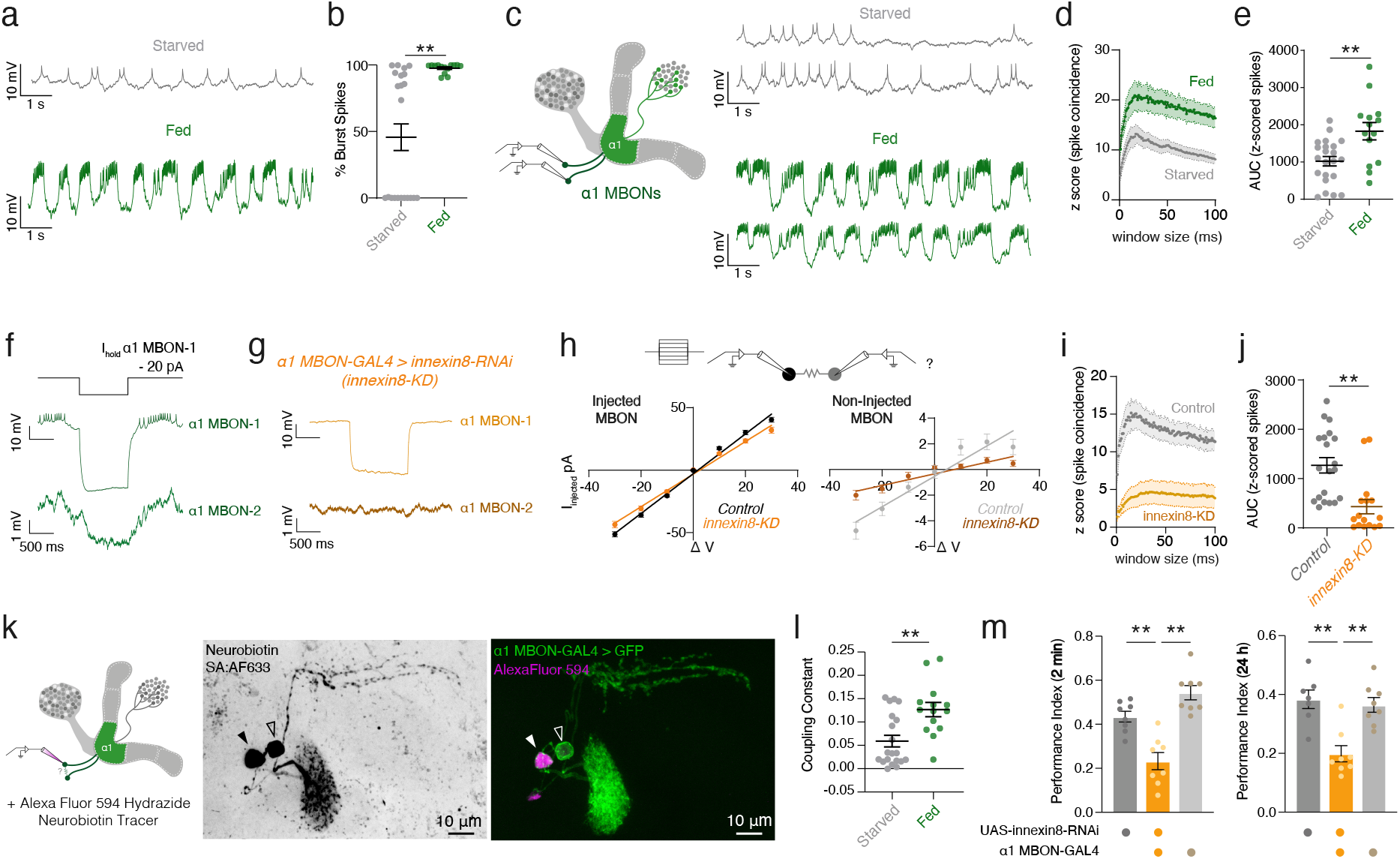
Hunger state regulates bursting and electrical coupling between a1 MBONs. **a,** Representative traces of spontaneous activity recordings of individual α1 MBON from starved and fed flies. **b,** Fed flies showed a higher percentage of burst spikes than starved (n=22, 14 neurons – 2 per fly; Mann-Whitney test, p<0.0001). **c,** Representative dual recordings in starved and fed flies. **d,** Spike coincidence (z-scored) versus detection window (n=22 starved, n=14 fed individual neurons – 2 per fly; mean ± s.e.m.). **e,** AUC for z-scored spike coincidence from **d** (Welch’s test, p=0.0064). **f,** Current injection into one α1 MBON (top, MBON1) elicits a voltage change in the other α1 MBON (bottom, MBON2). **g,** *innexin8* knockdown (*MB310C>innexin8-RNAi*) reduces MBON coupling; current injection into MBON1 no longer produces a change in MBON2. **h,** Current-voltage (I-V) plots for control and *innexin8* knockdown (*innexin8-*KD) pairs (Injected MBON; n=12 neurons – 2 per fly, *Control* R^2^=0.9664, p<0.0001; *innexin8-*KD R^2^=0.9284, p<0.0001). Non-injected MBON showed reduced coupling with *innexin8-*KD (n=16 neurons – 2 per fly, *Control* R^2^=0.5814, p<0.0001; *innexin8-*KD R^2^=0.4137, p<0.0001). **i,** Spike coincidence across detection windows, control (grey, n=20 neurons – 2 per fly) and *innexin8-KD* (orange, n=16 neurons – 2 per fly); mean ± s.e.m. **j,** AUC for **i** (Mann-Whitney test; p=0.0002). **k,** Left, neurobiotin injection into one α1 MBON. Middle, neurobiotin-labelled neurons (SA:AF633). Right, patched cell co-filled with non-gap-junction permeable AlexaFluor-594 Hydrazide. **l,** Coupling constant (ΔV_Cell2_/ΔV_Cell1_) in fed vs. starved flies (n=20,14 neurons – 2 per fly; Mann-Whitney test, p=0.0068). **m,** *innexin8-*KD in α1 MBONs reduces short- and long-term memory (One-way ANOVA with Dunnett’s multiple comparisons tests, STM; p=0.0005, p<0.0001, n=8,8,8 groups of flies. LTM; p=0.0003, p=0.0006, n=7,8,8 groups of flies).

We next investigated whether α1 MBON synchrony arises from gap junction coupling. Consistent with this possibility, current injection into one neuron produced a voltage deflection in its partner (Fig. 3f). To further test for electrical coupling, we knocked down *innexin8* (*inx8*), which encodes the principal gap junction subunit expressed in the *Drosophila* brain, in α1 MBONs ^32,33^. *Innexin8* knockdown in α1 MBONs reduced electrical coupling and coincident spiking between α1 MBONs relative to controls (Fig. 3g-j; Supplementary Fig. 3e,f). Finally, we performed dye-coupling experiments. Filling a single α1 MBON with gap-junction-permeable neurobiotin also labelled the other ipsilateral α1 MBON (Fig. 3k; Supplementary Fig. 3g).

Since *innexin8* knockdown reduced spike synchrony (Fig. 3i,j), we hypothesized that hunger-state-dependent modulation of gap junction coupling between α1 MBONs could explain differences in spike synchrony observed in starved and fed animals (Fig. 3d,e). Strikingly, quantification of coupling constants, defined as the ratio of the voltage deflection in the non-injected cell to that produced by current injection in its partner, revealed fed animals to have increased electrical coupling between α1 MBONs (Fig. 3l). Coupling strength also correlated significantly with z-scored coincident spike probability (Supplementary Fig. 3h), reinforcing the idea that gap junction coupling facilitates synchronization of spiking across ipsilateral α1 MBONs. Furthermore, loss of innexin8 suppressed the synchronized bursting mode characteristic of fed animals, leaving α1 MBONs in the tonic spiking mode (Supplementary Fig. 3l,m). Finally, *innexin8* knockdown in α1 MBONs impaired sucrose-reinforced short- and long-term olfactory memory (Fig. 3m). Importantly, *innexin8* knockdown did not alter resting membrane potential or spike amplitude, indicating that behavioral deficits did not merely result from silencing MBON excitability or broad physiological disruption (Fig. 3h; Supplementary Fig. 3i-k). Further, defective short-term memory suggests that gap junction-mediated synchronized spiking contributes to memory formation, consistent with evidence that the molecular and circuit processes underlying long-term memory are initiated during acquisition rather than emerging only during later consolidation ^34–35^.

### α1 circuit encoding ability depends on hunger state

Given variation in activity across DANs and MBONs, we investigated whether different physiological modes support distinct circuit-level operations. We therefore compared the effect of optogenetic α1 DAN stimulation on α1 MBON activity in starved versus fed animals. In starved animals, dopaminergic stimulation produced a robust increase in spiking in α1 MBONs, whereas the same stimulation had little to no influence in fed animals (Fig. 4a; Supplementary Fig. 4a). To determine if loss of dopamine responsiveness was due to a loss of ability for the postsynaptic neuron to respond (e.g. receptor internalization) or due to saturation of activity from elevated bursting characteristic of the fed state^36,37^, we hyperpolarized α1 MBONs to −60 mV by injecting a small holding current (<20 pA) to suppress baseline bursting activity. Hyperpolarizing MBONs was sufficient to restore a depolarizing response to dopamine neuron stimulation in fed animals (Supplementary Fig. 4b,c).

**Figure 4:**
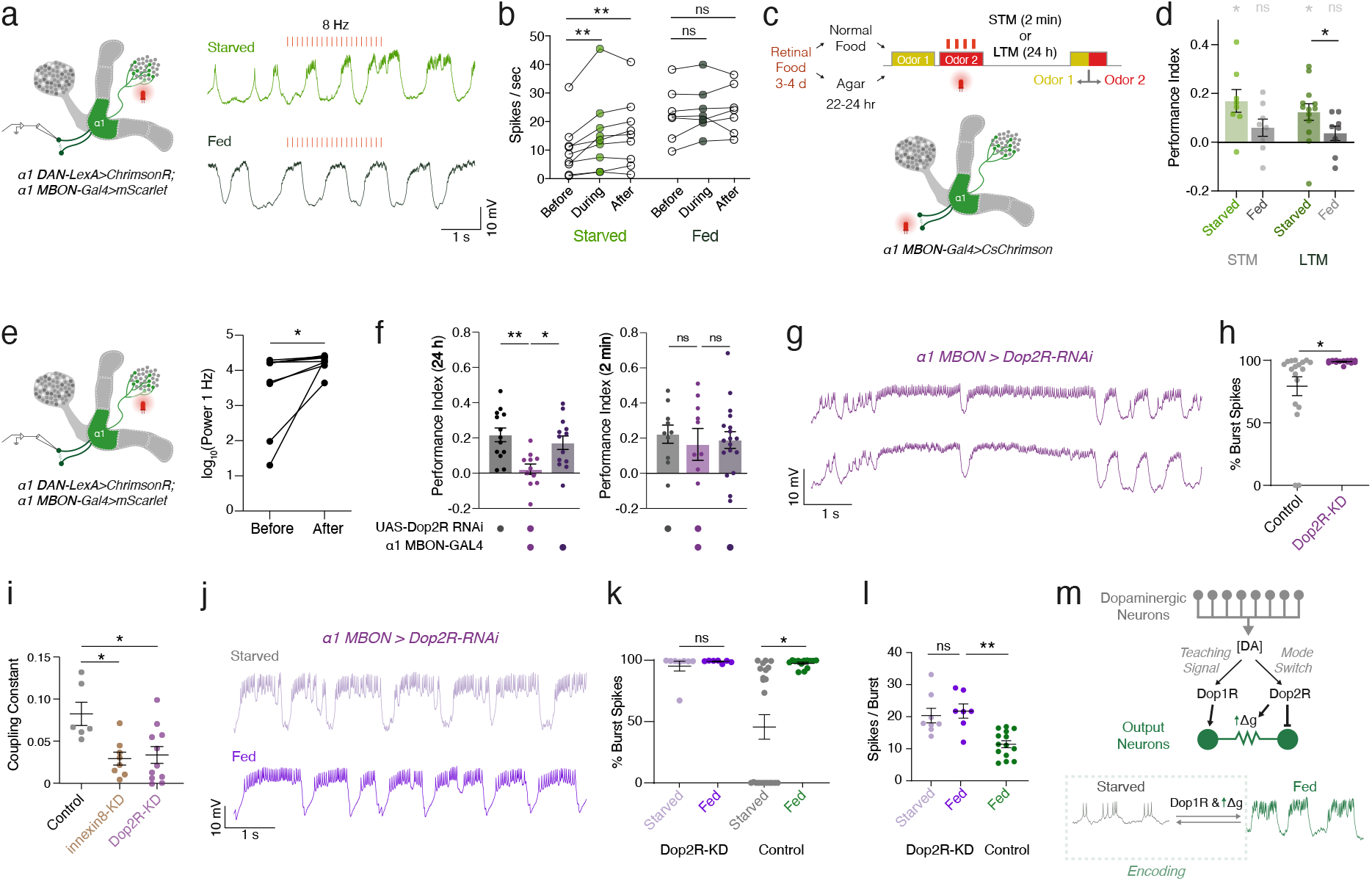
Internal state gates learning through gain modulation of a1 MBON activity. **a,** α1 MBON recordings during optogenetic α1 DAN stimulation in starved or fed flies. **b,** Spike frequency increases in starved (Friedman test; n=9 neurons – 1 per fly with repeated measures, Before vs. During p=0.0094, Before vs. After p=0.0008), not fed flies (repeated-measures one-way ANOVA with Geisser-Greenhouse correction, F(1.414,8.487)=0.9859, p=0.3815, ε=0.7072, partial η²=0.141; n=7 neurons – 1 per fly, Before vs. During p=0.1819, Before vs. After p=0.5279). **c,** Starved or fed flies were trained with α1 MBON light-stimulation paired with an odor then tested for odor preference. **d,** Starved flies preferred the light-paired odor (green) (One-sample t-test, µ=0, STM n=8 groups of flies, p=0.0084, LTM n=13 groups of flies, p=0.0034) but not fed flies (grey) (One-sample t-test, µ=0, STM n=8 groups of flies, p=0.1361, LTM n=8 groups of flies, p=0.2591). LTM was greater in starved vs. fed flies (Mann-Whitney, p=0.0351). **e,** 1 Hz power of α1 MBONs increases following 20 Hz stimulation of α1 DANs (Wilcoxon test; n=9 neurons – 1 per fly, p=0.0039). **f,** *Dop2R* knockdown (*Dop2R-*KD, purple) suppresses LTM in α1 MBONs (left, one-way ANOVA with Tukey’s multiple comparisons; n=13,11,13 groups of flies, UAS vs. *Dop2R*-KD p=0.0021, GAL4 vs. *Dop2R*-KD p=0.0194) but not STM performance (right, Kruskal-Wallis with Dunn’s multiple comparisons; n=10,10,18 groups of flies, p>0.05). **g,** Dual *Dop2R*-KD recording (compare to Fig. 3a,c). **h,** Burst-spike proportion is greater in *Dop2R*-KD vs. controls (Mann-Whitney, n=16,18 neurons – 2 per fly, p=0.0034). **i,** Coupling is reduced in *Dop2R-KD* vs. controls, like *innexin8-*KD (Kruskal-Wallis; n=6,8,11 neuron pairs, one per brain, Control vs. *innexin8-*KD p=0.0435, Control vs. *Dop2R*-KD p=0.0163). **j,** Representative traces of *Dop2R*-KD α1 MBONs in starved and fed flies. **k,** *Dop2R*-KD locks α1 MBONs in the bursting mode (Kruskal-Wallis; n=8,7,22,14 neurons (left to right), *Dop2R*-KD p>0.999, Control p=0.0005). **l,** Spikes-per-burst is greater in fed *Dop2R*-KD flies vs. controls (Kruskal-Wallis; n=8,7,14 neurons (left to right), *Dop2R*-KD Starved vs. Fed p>0.999, *Dop2R*-KD Fed vs. Control Fed p=0.0008). **m,** Dopamine’s net effect on α1 MBON optogenetic activation is stimulatory when flies are starved. Dopamine simultaneously acts on Dop2R, enhancing electrical coupling and suppressing bursting.

These results demonstrate that starvation puts the α1 DANs and MBONs into a low-activity mode, which may be a prerequisite for memory formation. To test whether such a transition from a low- to high-activity mode is required for memory formation, we abruptly induced a high-activity mode by optogenetically activating α1 MBONs coincident with odor presentation in low-activity-mode starved flies (Fig. 4c). Strikingly, starved flies showed short- and long-term memory for the paired odor, whereas sated flies did not (Fig. 4d), indicating that the transition into the high-activity mode, and not the high-activity mode alone, is what drives memory formation.

### Dopamine drives network mode transitions

We next investigated whether α1 DAN activity switches baseline activity of α1 MBONs into the bursting physiological mode representing satiation. Optogenetic stimulation of α1 DANs increased bursting in α1 MBONs, consistent with α1 DANs driving network transitions (Fig. 4e). To identify the memory-relevant dopamine receptors in α1 MBONs, we knocked down their expression using RNAi ^38^. Knockdown of *Dop2R* abolished long-term sucrose-reinforced memory but not short-term memory, whereas knockdown of *Dop1R* or *DopEcR* had no effect (Fig. 4f; Supplementary Fig. 4d,e). We therefore examined the physiological consequences of *Dop2R* knockdown in α1 MBONs. *Dop2R* knockdown produced an increase in burst spiking (Fig. 4g,h; Supplementary Fig. 4f) but did not change total spike frequency (tonic and burst spiking) or membrane potential (Supplementary Fig. 4g,h).

Surprisingly, gap junction coupling between α1 MBONs was also reduced in *Dop2R* knockdown animals (Fig. 4i; Supplementary Fig. 4j). Taken together, Dop2R specifically modulates burst spiking and gap junction coupling between α1 MBONs.

We noted that *Dop2R* knockdown elevated bursting above fed controls, raising the possibility that loss of Dop2R signalling locks α1 MBONs in the bursting mode representing the fed state. In support of this idea, *Dop2R* knockdown locked α1 MBONs into a bursting mode irrespective of satiation state (Fig. 4j,k; Supplementary Fig. 4k,l), and elevated bursting and membrane potential above that seen in fed genetic control flies (Fig. 4l; Supplementary Fig. 4m).

To briefly summarize, these findings demonstrate that dopamine serves complementary functions through opposing receptor mechanisms. Reward-evoked dopamine can increase α1 MBON activity only if flies are hungry prior to training (Fig. 4a,b). Dopamine simultaneously transitions the network into a satiated mode by engaging Dop2R-dependent enhancement of gap junction coupling between MBONs and constraining burst intensity (Fig. 4m). We propose that this satiated mode represents how strongly the circuit commits the network to consolidating a recently acquired memory. Dop2R limits this commitment by dampening burst intensity while still preserving synchrony through gap junction coupling.

Such a mechanism could maintain newly formed appetitive memories in a transiently labile state, susceptible to delayed post-ingestive information to either promote or suppress consolidation depending on whether the consumed food is nutritious or toxic ^12,25,39–42^.

### Activity modes are coordinated across α1 DANs and MBONs

Our state-transition model suggests that activity modes of DANs and MBONs could be coordinated. We therefore performed dual whole-cell patch-clamp recordings from α1 DAN– MBON pairs (Fig. 5a) to relate DAN spike modes to the bursting mode of its partner MBON in the same fly. Because α1 DANs comprise three morphological subtypes and it is unknown if they have different physiological properties or functional post-synaptic targets (e.g. excitatory or inhibitory dopamine receptors), we filled recorded DANs with biocytin for post-hoc morphological identification (Fig. 5b,c; Supplementary Fig. 5a), allowing subtype assignment in 16 out of 23 pairs. Critically, flies were ad libitum fed on standard food prior to recording to preserve natural variation in hunger state, which revealed substantial variability in spontaneous activity across DANs and MBONs. Strikingly, DAN spiking pattern was tightly correlated with MBON activity mode, with large-amplitude spiking DANs associated with bursting MBONs (Fig. 5d; Supplementary Fig. 5b). These results demonstrate that physiological modes are largely coordinated across the DAN–MBON circuit.

**Figure 5:**
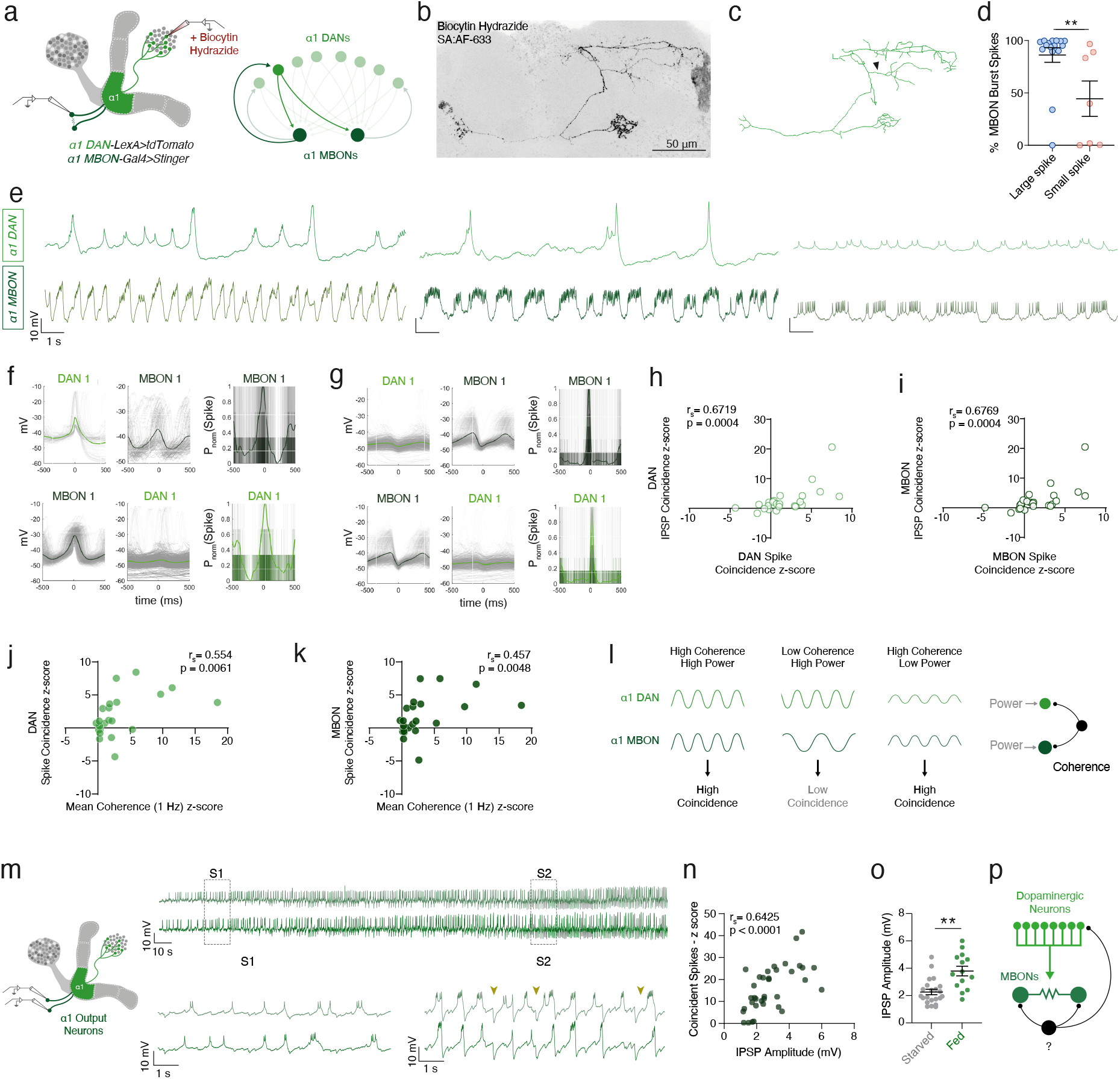
Coherence differentiates physiological states of the a1 compartment. **a,** Schematic of dual α1 DAN and MBON recordings (only α1 DANs were biocytin filled). **b,** Representative image of biocytin filled α1 DAN. **c,** Example reconstruction of **b**. **d,** MBONs paired with large-amplitude-spiking DANs show greater burst spiking compared to small-amplitude-spiking DANs (n=16,7 neurons – 1 MBON per fly; Mann-Whitney test two-tailed, p=0.0076). **e,** Traces from three dual recordings from different flies, representing variation in natural hunger state. Top, DANs; bottom, MBONs. **f,** Representative spike-triggered average for a coherent, coincidentally spiking DAN-MBON pair. **g,** IPSP-triggered average for the same pair. **h,i,** DAN and MBON spike coincidence (z-scored) correlate with IPSP coincidence (z-scored) (Left, Spearman correlation; n=23 flies – one DAN-MBON pair per fly, p=0.0004, r=0.6719. Right, Spearman correlation; n=23 flies – one DAN-MBON pair per fly, p=0.0004, r=0.6769). **j,** Mean coherence between DAN-MBON pairs correlates with DAN spike coincidence (z-scored) (Spearman correlation; n= 23 flies – one DAN-MBON pair per fly, p=0.0061, r=0.5543) and with **k,** MBON spike coincidence (z-scored) (Spearman correlation; n=23 flies – one DAN-MBON pair per fly, p=0.0048, r=0.5672). **l,** α1 DAN and MBON coherence can vary independently of power to shape coincidence. 1 Hz coherence could be generated by a common inhibitory input (black) onto α1 DANs and MBONs, that entrains their 1 Hz oscillations. **m,** Dual α1 MBON recordings occasionally showed spontaneous transition states. Example trace in which a transition occurs from state 1 (S1) to state 2 (S2). Large IPSP amplitudes are a feature of S2 (yellow arrowheads). **n,** IPSP amplitude correlates with coincident spikes (z-scored) (Spearman correlation; n=36 neurons – two MBONs per fly, r=0.6425, p<0.0001). **o,** IPSP amplitude increases in fed flies vs. starved (Mann-Whitney test; n=14,22 neurons – two MBONs per fly, p=0.0006). **p,** Schematic of putative origins of enhanced IPSP amplitude (black neuron).

Across our recordings, DAN-MBON pairs varied in their spiking modes – likely representing different relative hunger states (Fig. 5e) – and also temporal coordination of spiking (Fig. 5f,g). Such coordination could govern circuit function through mechanisms such as spike-timing-dependent plasticity or shared phase relationships that permit certain circuit elements to communicate ^43,44^. To examine how coordination emerges in the α1 circuit, we therefore performed a more detailed analysis of physiological properties. Specifically, we examined coincidence of fast events such as spiking and inhibitory post-synaptic potentials (IPSPs) as well as coherence and power at 1 Hz. We explicitly examined the 1 Hz frequency range because both the α1 DAN and MBON power spectra showed a peak in power around 0.5–1.5 Hz (Supplementary Fig. 3a, Supplementary Fig. 5c-f).

From our analyses, we observed that DAN–MBON spike coincidence varied across recordings (Fig. 5f,g; Supplementary Fig. 5g). However, spiking mode alone was not predictive of synchronized spiking across DAN-MBON pairs (e.g. large-amplitude-spiking dopaminergic neurons were not always coincidentally spiking with bursting MBONs). As a result, 1 Hz oscillatory power of individual cells was not predictive of DAN-MBON coincident spiking or IPSPs (Supplementary Fig. 5h,i). These findings suggest that a separate mechanism underlies spike mode generation and coordination across the circuit.

Spike coincidence, however, was tightly correlated with the degree of coincident IPSPs (Fig. 5h,i), suggesting that a common inhibitory input could coordinate spiking across DANs and MBONs by constraining their spike timing. Additionally, spike and IPSP coincidence correlated with slow 1 Hz coherence between DAN and MBON pairs (Fig. 5j,k; Supplementary Fig. 5j,k), demonstrating that coordination of slower frequencies helps align fast events. Taken together, these results indicate that precise temporal coordination depends on slow coherence between neurons and not simply spiking mode (Fig. 5l). Finally, to determine if differences in spiking mode and coincident activity reflect functional heterogeneity of anatomical α1 DAN subtypes, we performed a principal component analysis (PCA) across physiological features (Table 1). The anatomical subtypes distributed evenly across PC space, indicating that all α1 DAN subtypes have the capacity to produce large- and small-amplitude spikes and vary in their coincident activity (Supplementary Fig. 5l-n).

Given our observed hunger-state-dependent differences in MBON spiking synchrony, we also examined whether inhibition could synchronize ipsilateral α1 MBON pairs. Intriguingly, in some dual α1 MBON recordings, spontaneous transitions between bursting modes were accompanied by changes in IPSP amplitude (Fig. 5m). When we examined IPSP amplitude against the degree of coincident spiking across dual recordings, we observed a strong positive correlation (Fig. 5n), demonstrating that stronger inhibition sharpens the temporal precision of coincident spiking. Finally, IPSP coincidence and amplitude, but not rate, were elevated in α1 MBON pairs in fed animals (Fig. 5o; Supplementary Fig. 3n,5o), indicating that the magnitude of inhibitory drive onto the α1 MBONs is itself state-regulated.

The relationship between IPSP amplitude and spike coincidence was specific to MBON-MBON pairs. Spike coincidence of DAN-MBON pairs only correlated with IPSP coincidence rather than IPSP amplitude (Fig. 5h,i), revealing that coincident inhibition, rather than its strength, coordinates DAN-MBON spiking. Together, these results reveal two means by which inhibition coordinates temporal activity within the α1 circuit: shared inhibitory timing aligns spiking across DAN-MBON pairs, whereas inhibitory gain sharpens coincident spiking between ipsilateral MBONs.

### Rhythmic inhibition instructs memory consolidation

We next sought to identify the source of inhibitory input to the α1 circuit and determine the function of the coherent activity it entrains. Prior work has described a motif of inhibitory MBONs (GABAergic γ1-pedc MBON and glutamatergic β1>α MBON) that interact with the α1 DAN-MBON circuit ^13^, some of which have been demonstrated to have roles in appetitive memory consolidation. In addition, odor-evoked activity of γ1-pedc MBON has been shown to be hunger-modulated, which is important for appetitive memory consolidation and expression ^23,39,45^. However, to date, no study has demonstrated a functional role for this inhibitory MBON motif in memory-relevant functions of the α1 circuit. On revisiting the synaptic connectivity of the inhibitory MBON motif, we found that α1 DANs are also part of this network (Fig. 6a). Therefore, the inhibitory MBON circuit motif provides a parsimonious explanation for the synchronized IPSPs we observed between α1 DANs and MBONs (Fig. 5j-l).

**Figure 6:**
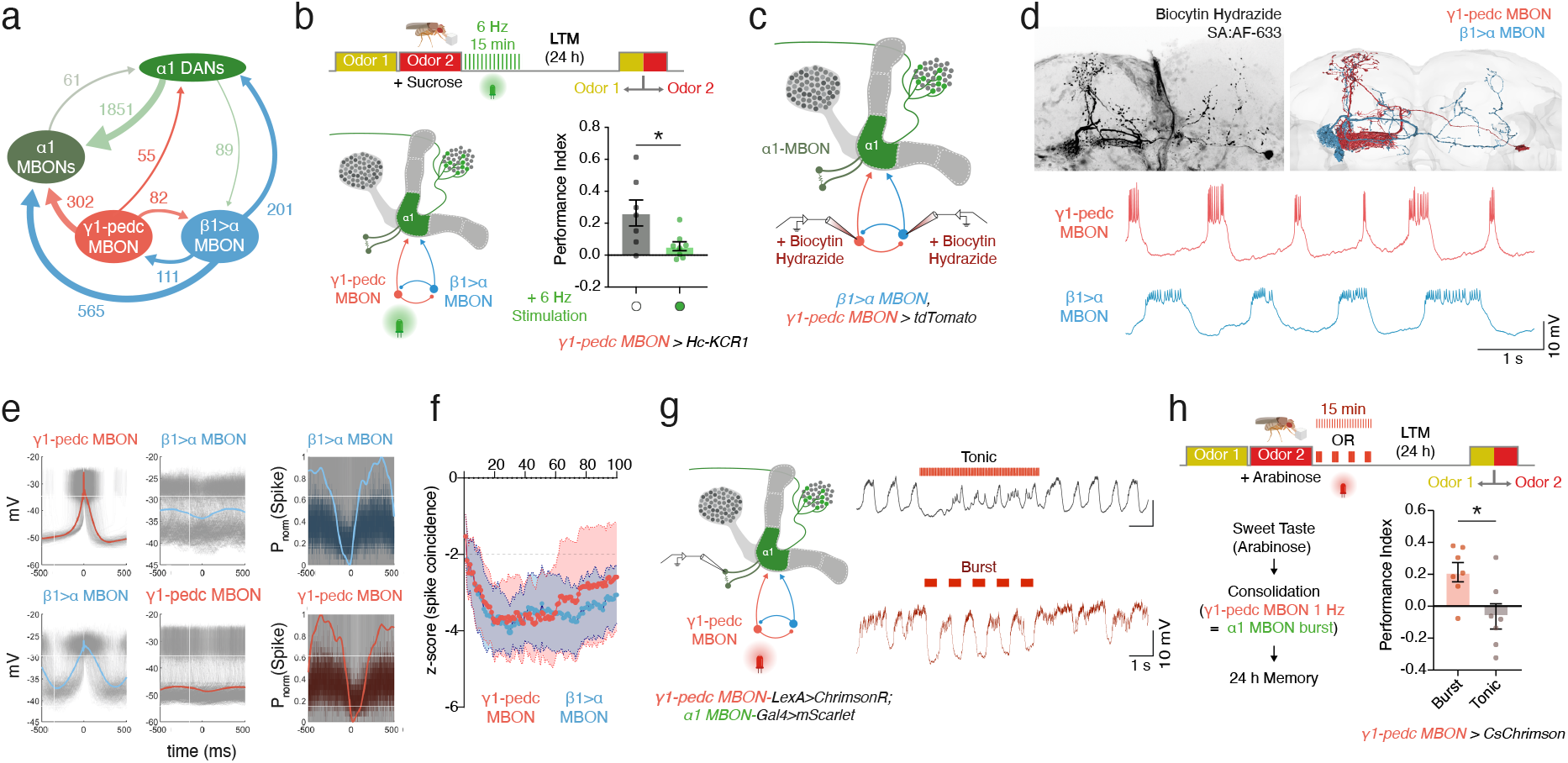
Rhythmic inhibition from Y1-pedc MBON stabilizes bursting in a1 MBONs and supports consolidation. **a,** Connectome-based circuit schematic of the α1 DAN, α1 MBON, Y1-pedc MBON, and β1>α MBON. Numbers indicate the total number of synapses per cell-type (FlyWire FAFB). **b,** Optogenetic inhibition of Y1-pedc MBON for 15 min immediately after sucrose training (6 Hz, 5 ms pulse width) decreased long-term memory performance (Welch’s test; n=7,8 groups of flies, p=0.0440). **c,** Schematic for dual recordings of Y1-pedc MBON and β1>α MBON; both neurons were filled with biocytin hydrazide. **d,** Top left, representative images of biocytin labelled neurons. Top right, EM reconstruction of a Y1-pedc MBON and β1>α MBON (FlyWire FAFB). Bottom, representative trace of a dual recording from a Y1-pedc MBON and β1>α MBON pair. **e,** Spike-triggered averages of an example Y1-pedc MBON and β1>α MBON pair and the normalized spike probability. Y1-pedc MBON and β1>α MBONs show anti-correlated spike timing. **f,** Z-scored spike coincidence between Y1-pedc and β1>α MBON pairs shows anti-correlated spiking (n=5 pairs, one pair per brain; mean ± s.e.m.). Values below zero indicate coincidence lower than shuffled values. Dotted horizontal line indicates z=-2 significance threshold. **g,** Representative traces of α1 MBONs with optogenetic stimulation of Y1-pedc MBON in a pattern that elicits tonic spiking (top trace) or burst spiking (bottom trace). **h,** Flies trained with non-nutritive arabinose were then exposed to optogenetic stimulation of Y1-pedc MBONs immediately following training for 15 min, then tested for memory 24h after training. Flies that had optogenetic stimulation that elicited burst firing in α1 MBONs exhibited an appetitive memory, whereas tonic stimulation did not (Welch’s test; n=7 groups of flies, p=0.0176).

Given that the high-activity bursting mode occurs following sugar consumption and is characteristic of a satiated state, we hypothesized that bursting activity in α1 MBONs immediately after feeding could play an active role in memory consolidation. We therefore tested whether γ1-pedc MBONs are required immediately after training, by silencing them for 15 min using expression of the light-gated potassium channel HcKCR^46^. Transient post-training silencing of γ1-pedc MBONs impaired sucrose-reinforced olfactory long-term memory (Fig. 6b; Supplementary Fig. 6a), demonstrating an early requirement for γ1-pedc MBONs in memory consolidation.

Given the importance of γ1-pedc MBONs during consolidation, we tested whether the inhibitory MBON circuit could drive the bursting mode of α1 MBONs. We reasoned that the γ1-pedc MBON and β1>α MBON circuit could form an antiphase oscillator, to entrain the 1 Hz rhythmic bursting of α1 MBONs. To investigate this, we performed dual patch-clamp recordings from pairs of γ1-pedc MBON and β1>α MBONs to examine their functional interaction. Because the MB434B driver line labels β1>α MBONs and γ4>γ1γ2 MBONs, unambiguous identification of recorded cell types required post-hoc verification. We therefore filled both pipettes with biocytin to allow for morphological identification of recorded neurons (Fig. 6c,d; Supplementary Fig. 6b). We observed anti-correlated spiking in five confirmed γ1-pedc MBON and β1>α MBON pairs (Fig. 6e,f; Supplementary Fig. 6c-e).

Consistent with their reciprocal connectivity, depolarizing current injection into β1>α MBON produced inhibition in γ1-pedc MBON, whereas the reciprocal manipulation produced a non-significant reduction of β1>α MBON voltage (Supplementary Fig. 6f). Although these data suggest that these MBONs are asymmetrically connected, theoretical and experimental studies have shown that stable antiphase activity can be sustained by weak, asymmetric reciprocal inhibition, particularly in neurons with post-inhibitory rebound ^47–50^. Indeed, we observed post-inhibitory rebound with hyperpolarizing current injection in both γ1-pedc MBON and β1>α MBON (Supplementary Fig. 6g). We therefore propose that the β1>α and γ1-pedc MBON motif provides rhythmic inhibition to the α1 circuit (see Supplementary Fig. 6h).

Finally, to determine whether rhythmic bursting in the α1 MBONs was sufficient to drive consolidation of long-term memories, we optogenetically stimulated γ1-pedc MBONs to evoke a tonic or bursting mode in α1 MBONs (Fig. 6g) following arabinose training. We applied these stimulation patterns for 15 min immediately after training with arabinose, a sweet non-nutritive sugar, which reinforces short-but not long-term memory ^7^. Remarkably, burst-patterned stimulation was sufficient to convert arabinose experience into long-term memory, whereas tonic stimulation did not (Fig. 6h). Therefore, memory consolidation emerges from two interdependent mechanisms, persistent large-amplitude spiking in α1 DANs and temporally structured inhibition of the α1 DAN-MBON circuit. Together, these two mechanisms establish a coherent activity mode, which transitions the circuit out of the learning-competent mode to one that consolidates recently formed memories.

## Discussion

We discovered that hunger state reconfigures the α1 circuit between two physiological modes that differ computationally, algorithmically, and in their molecular implementation. At the computational level, the circuit engages a learning-competent mode when flies are hungry; once satiated, it transitions into a mode that consolidates recently formed memories, and if satiation is complete, this mode also prevents further learning.

At the algorithmic level, the mode transition reflects a shift from a low-to a high-activity reward state. In hungry animals, low coherence among circuit elements prevents α1 DANs from integrating inputs, and MBONs remain in a tonic activity mode. When a food-related reward signal robustly activates α1 DANs, the circuit rapidly transitions into a coherent, high-activity mode; memory encoding occurs specifically as a consequence of this low-to-high transition. Following satiation, the circuit remains locked in this coherent high-activity mode, in which persistent large-amplitude DAN spiking drives sustained MBON bursting and reduces the gain of further reward signals on MBON activity.

At the implementation level, we identify molecules that establish and stabilise each mode. During starvation, dNPF-mediated signalling suppresses large-amplitude spiking in α1 DANs, establishing the low baseline reward state described above. During food ingestion, activation of sweet-taste receptor neurons drives large-amplitude DAN spiking. This increase in dopaminergic activity depolarises α1 MBONs and transitions them from tonic to burst firing (i.e. low-to-high-activity transition). This post-feeding, high-activity MBON mode is stabilised by Dop2R-dependent regulation of bursting through enhanced gap junction coupling, together with rhythmic inhibitory input from the γ1-pedc MBON. Rhythmic inhibition enhances coherence across the α1 circuit (DAN to MBON) and imposes the bursting mode in MBONs.

A central unexpected finding was that α1 DANs generate large-amplitude spikes through coincident activation of multiple spike initiation zones. Our model provides an intriguing possibility that the coincident SIZ activation mechanism could be important for detecting simultaneous inputs from the dendrites and axons^51,52^. During sugar-reinforced olfactory training, flies feeding on sucrose receive excitatory dendritic inputs into SIZ 3, which provide representations of reward information (e.g. sweet taste) and simultaneous sensory-evoked axo-axonal inputs from cholinergic Kenyon Cell activation, which could generate back-propagating action potentials from SIZ 2^21,22^. The convergence of these two signals may therefore promote large-amplitude spiking and provide an instructive signal for learning. This mechanism has interesting similarities to cerebellar Purkinje cell complex spikes, as both systems use compartmentalized excitability to generate specialized spike outputs associated with instructive signals for plasticity ^53^. Whether large- and small-amplitude spikes are functionally distinguished at the level of dopamine released remains unresolved. Our biophysical model suggests they are not obviously differentiated at axon terminals, though boutons could still be sensitive to spike waveform and produce distinct modes of transmitter release ^54–56^. Alternatively, large-amplitude spikes may act within the soma itself – e.g. activation of activity-dependent transcription factors such as CREB, whose nuclear localisation and transcriptional activity are sensitive to spike amplitude and pattern ^57^.

The state-dependent switch in α1 MBON activity may have broader implications for general MBON function. MBONs receive convergent input from thousands of Kenyon cells and project to diverse downstream targets regulating approach and avoidance behaviors.

Changes in MBON firing mode alter not only the output strength but also the effective connectivity of downstream circuits. This resembles other systems in which firing mode dictates circuit function, such as thalamic neurons switching between tonic and burst modes to reorganize thalamocortical communication, or stomatogastric neurons switching oscillatory regimes to participate in different network rhythms ^48,58,59^. The hunger-state-dependent switching we observe in α1 MBONs may therefore configure downstream circuit function independently of odor-evoked memory function^28^. More broadly, because MBONs are highly interconnected and converge onto shared downstream targets across different MBON types^13^, the MBON layer may operate as a state-dependent switchboard in which oscillatory coherence across different MBON types determines which outputs downstream of the MBONs are functionally coupled at a given time.

We identify a synchronized activity mode across DANs and MBONs that is organized by inhibitory inputs from the β1>α and γ1-pedc MBONs. This synchrony could gate plasticity between coherently active circuit elements ^60,61^, accounting for why this mode may be important for memory consolidation. Post-feeding alignment of DAN-MBON activity may bias recently active synapses towards consolidation^62^. This is reminiscent of consolidation following contextual fear conditioning in hippocampal area CA1, where principal cells increase their coherence after training through enhanced gain of inhibitory parvalbumin-expressing interneurons^63^. In CA1, this coherence is closely associated with replay of previously active neuronal ensembles, a process thought to drive consolidation^64^.

**Table 2:** Model parameters.

| Parameter | Value | Description |
| --- | --- | --- |
| $E_L, E_{Na}, E_K, E_{Ca}, E_{CAN}$ | -40, +50, -77, +120, 0 mV | Reversal potentials |
| $\bar{g}_L, \bar{g}_{Na}, \bar{g}_K, \bar{g}_{Ca}, \bar{g}_{SK}, \bar{g}_{CAN}$ | 1.4, 80, 50, 0.1, 8, 12 mS cm<sup>-2</sup> | Maximum conductance densities |
| $C_m$ | 10 μF cm<sup>-2</sup> | Specific membrane capacitance |
| $R_i$ | 25 Ω cm | Axial resistivity |
| $m$ (Na act.) | $V_{1/2} = -23$ mV, $a = 0.085$ mV<sup>-1</sup>, $\tau = 40$ ms | Sodium activation |
| $h$ (Na inact.) | $V_{1/2} = -30$ mV, $a = -0.05$ mV<sup>-1</sup>, $\tau = 60$ ms | Sodium inactivation |
| $n$ (K act.) | $V_{1/2} = -23$ mV, $a = 0.06$ mV<sup>-1</sup>, $\tau = 150$ ms | Potassium activation |
| $p$ (Ca act.) | $V_{1/2} = -20$ mV, $a = 0.1$ mV<sup>-1</sup>, $\tau = 100$ ms | Calcium activation |
| $s$ (SK) | $Ca_{1/2} = 0.02$ mM, $a = 400$ mM<sup>-1</sup>, $\tau = 500$ ms | SK gating |
| $r$ (CAN) | $Ca_{1/2} = 0.02$ mM, $a = 400$ mM<sup>-1</sup>, $\tau = 2000$ ms | CAN gating |
| $\tau_{Ca}, \alpha_{Ca}$ | 1000 ms, 0.005 mM m<sup>2</sup> ms<sup>-1</sup> A<sup>-1</sup> | Calcium pool |

## Acknowledgements

We thank J. Ache, M. Wojcik, T. Falt, J. Felsenberg, and A. Cook for helpful discussions and comments on the manuscript. We are grateful to Y. Aso, H. Amrein, Bloomington Drosophila Stock Center (NIH P40OD018537), Vienna Drosophila Resource Center and HHMI Janelia Research Campus for fly stocks. Y. Z. was funded by a Wellcome Early Career Award (226333) to A. P.. B. S., J. C., C. D. T. and A.P. were funded by a Wellcome Principal Research Fellowship (200846), a Wellcome Discovery Award (225192), an ERC Advanced Grant (789274), and Wellcome Collaborative Awards (203261 and 209235) to S. W.. N.M., S. S. B. and L.F. were funded by the Max Planck Society. I. C. and R. P. C. were funded by EPSRC (X029336) and ERC-UKRI Frontier Research Guarantee Grant (Y027841).

## Author contributions

Designed **research** A. P., N. M., J. C., I. C., S. W., Performed research A. P., B.S., Y.Z., J. C., N. M., I. C., C. D. T., Analyzed data A. P., B. S., J. C., Y. Z., N. M., I. C., C. D. T., Resources A. P., S. W., Writing A. P., S. W., Supervision A. P., S. W., L. F., S. B., Funding Acquisition A.P., S.W., R. P. C., L. F., S. B..

## Methods

### Drosophila strains

Driver Lines (GAL4):

MB043C-GAL4 (BDSC #68363)

MB310C-GAL4 (BDSC #68313) ^65^

MB112C-GAL4 (BDSC #68263) ^65^

MB434B-GAL4 (BDSC #68325)^65^

MB043C-LexA (58E02-ZpLexADBD in JK22C; 32D11-p65ADZp in JK73A) (Janelia – Aso Lab)

MB112C-LexA (SL86058 Split LexA R93D10 JK22C R13F04 VK00027 3040939) (Janelia – Aso Lab)

MB310C-LexA (SL86055 Split LexA R52G04 JK22C R17C11 VK00027) (Janelia – Aso Lab)

Gr5a^LEXA^ – Amrein Lab^66^ Gr43a^LexA^ – Amrein Lab^66^ Gr64f^LexA^ – Amrein Lab^66^

w[*]; TI{2A-lexA::GAD}NPF[2A-lexA]/TM3, Sb[1] (BDSC 84422)

Responder Lines (UAS):

w/w,13XLexAop2-IVS-ChrimsonR-mVenus-p10 attP18, 20XUAS-syn21 mScarlet-opt-p10 su(Hw)attp8 (From Yoshi Aso)

w; UAS-Flp, UAS-mCD8::GFP/CyO (From Wilson Lab, Harvard) y[1] w[*]; P{w[+mC]=UAS-FLP.D}JD1 (BDSC 4539)

#### Optogenetic Actuators

w[*]; P{y[+t7.7] w[+mC]=UAS-HcKCR1.ET.EYFP}attP40/CyO (BDSC 605298)

w[1118]; P{y[+t7.7] w[+mC]=13XLexAop2-IVS-CsChrimson.mVenus}attP40 (BDSC 55138) PBac{y[+mDint2] w[+mC]=13XLexAop2-IVS-CsChrimson.tdTomato}VK00005 (BDSC 82183)

#### RNAi Lines

Dop1R1 RNAi (VDRC v107058): P{KK102341}VIE-260B Dop2R RNAi (VDRC v11470 ): w1118; P{GD732}v11470/TM3 DopEcR RNAi (VDRC v103494): P{KK111211}VIE-260B npfr RNAi (VDRC v107663) : P{KK112704}VIE-260B npfr RNAi (VDRC v9605): w1118; P{GD668}v9605 shakB RNAi (VDRC v26801): w1118; P{GD12666}v26801

#### Fluorophores / Indicators

w[*]; P{w[+mC]=UAS-Stinger}2, PBac{y[+mDint2] w[+mC]=13XLexAop2-IVS-tdTomato.nls}VK00022 (BDSC 66680)

y[1] w[67c23] para[MI08578-mCherry.0] Mi{PT-mCh.0}MI08578a Mi{PT-mCh.0}MI08578b (BDSC 91529)

y[1] w[*] Mi{FlpTag.0}para[MI08578-FT.0-X] (BDSC 92146)

w[*]; P{y[+t7.7] w[+mC]=10XUAS-IVS-myr::tdTomato}attP2 (BDSC 32221) w[*]; P{y[+t7.7] w[+mC]=10XUAS-IVS-myr::tdTomato}attP40 (BDSC 32222) w[*]; P{y[+t7.7] w[+mC]=10XUAS-IVS-myr::GFP}attP40 (BDSC 32198) w[*]; P{y[+t7.7] w[+mC]=10XUAS-IVS-myr::GFP}attP2 (BDSC 32197)

w[1118]; P{y[+t7.7] w[+mC]=20XUAS-IVS-jGCaMP7f}su(Hw)attP5 (BDSC 80906)

### Behavior

#### Food deprivation

Flies were aliquoted into groups of ∼100 before behavioral experiments. For starvation, flies were food-deprived for 20 to 24 h in a 25 ml vial containing a 2 cm × 3 cm piece of filter paper with 1% agar at the base. Vials were stored at 23-25 °C throughout the starvation period.

#### T-maze olfactory behavioral experiments

Virgin female flies from GAL4 driver lines were crossed with male flies carrying the corresponding effector transgenes. Mixed-sex progeny aged 4–12 days were used for T-maze behavioral experiments, with approximately 100 flies tested per group.

Olfactory conditioning was performed using a T-maze apparatus (CelExplorer Labs, TM-101). 4-methylcyclohexanol (MCH) and 3-octanol (OCT) were used as conditioned odors^67^ (Sigma-Aldrich). Each odor was diluted ∼1:10^3^, specifically, 10-12.9 µl MCH or 7-8 µl OCT in 8 ml mineral oil (Sigma-Aldrich). All experiments were performed at 23 °C and 55% to 65% relative humidity.

#### Olfactory conditioning with sucrose

For experiments in Fig. 1i, 3m, 4f, 6b, and Supplementary Fig. 1a, 1d, 4d, and 4e sucrose was prepared as a super-saturated solution (∼400 g l^−1^), of which 3 ml was pipetted onto a 5 cm × 8 cm piece of filter paper. Excess solution was drained from the paper, which was then rolled into a tube and allowed to dry overnight. Appetitive training with sucrose was conducted as previously described^68^. Groups of flies were first transferred to a training tube of a T-maze lined with filter paper and then exposed to an odor (the CS−) for 2 min.

Following 30 s of clean air in the same tube, flies were transferred to another tube lined with dried sucrose then immediately exposed to another odor (the CS+) for 2 min. For STM, trained flies were transferred to the T-maze elevator after CS+ exposure and immediately given a choice between the CS− and CS+ odors. For LTM, trained flies were transferred to agar vial for 24h until testing.

#### Olfactory conditioning with optogenetic activation

For experiments in Figs. 2d, 4d and 6h, olfactory conditioning combined with optogenetic activation was performed as previously described^69^. Groups of flies were transferred into a tube on which three red LEDs (700 mA, centred at 630 nm, 3 W maximum power; Multicomp) were mounted. Flies were first exposed to the CS− odor for 2 min, followed by 30 s of clean air, and were subsequently exposed to the CS+ odor for 2 min. During CS+ presentation, red-light stimulation was delivered at the indicated frequency (see below). Light pulses were controlled using a DG2A Train/Delay Generator (Digitimer) coupled to a DS2A Isolated Constant Voltage Stimulator (Digitimer). For experiments in Fig. 2d, high-frequency stimulation (HFS) consisted of 15 ms pulses delivered at 20 Hz for trains of 2.5 seconds for 2 minutes. Low-frequency stimulation (LFS) consisted of 15 ms pulses delivered at 2.92 Hz continuously for 2 minutes. For short-term memory (STM) experiments, flies were tested for their choice between the CS− and CS+ odors following training. For long-term memory (LTM) experiments, trained flies were transferred to vials containing 1% agar and maintained horizontally in a cardboard box until memory was tested 24 h after training.

For the experiments in Fig. 4d, neural activation was induced using 5 ms pulses delivered at 10 Hz in trains of 1 sec with an inter-train-interval of 1 sec. Following training, flies assigned to the fed condition were transferred to yellow food vials, whereas flies assigned to the starved condition were transferred to vials containing 1% agar. Memory was subsequently tested immediately (STM) or 24 h (LTM) after training.

For the experiments in Fig. 6h, flies were conditioned using non-nutritive arabinose. Arabinose solution (3M) was applied to filter paper and allowed to dry before training. Immediately following training, flies received optogenetic stimulation for 15 min. To induce burst activity, stimulation consisted of 500 ms long continuous light pulses with a 500 ms inter-pulse-interval. To induce tonic activity, stimulation consisted of 15 ms pulses delivered at 20 Hz for 15 minutes. Memory was tested 24 h after training.

#### Olfactory conditioning with optogenetic inhibition

For the experiment in Fig. 6b, olfactory conditioning with sucrose combined with optogenetic inhibition was performed as described above for optogenetic activation, with the following modifications. Groups of flies were transferred to a training tube equipped with three green LEDs (700 mA, peak wavelength 530 nm; Multicomp). Flies were exposed to the CS− odor for 2 min, followed by 30 s of clean air, and were then transferred to a second tube for a 2-min exposure to the CS+ odor. Following training, flies received optogenetic inhibition for 15 min using 5 ms pulses delivered at 6 Hz for 15 minutes. Memory was tested 24 h after training.

#### Fly Husbandry

Flies were kept on standard cornmeal food under a 12:12 light:dark cycle at 60% humidity and 25°C unless otherwise stated. Flies used for electrophysiology were 1-3 days old. If flies were starved, they were kept on vials containing 1% agar for 18-22 hours prior to recordings. Flies refed sucrose prior to recording were refed with a Kimwipe soaked in 1 M sucrose solution in distilled water until satiation (when the fly did not accept more sucrose solution).

Flies used for optogenetics experiments were always kept in light-shielded boxes or foil-covered vials prior to the start of the experiment. Flies were raised on non-retinal containing standard cornmeal food and transferred to 1 mM retinal rich (all-*trans*-retinal) food after eclosion. Flies used for electrophysiology were transferred to retinal rich food for 2 days and recordings were performed when flies were 3-4 days old. Flies used for behavioral experiments were transferred to retinal rich food for 3 days and were 4-8 days old at the time of training.

### Immunohistochemistry / Staining

2-5 day old adult flies were anaesthetized with ice. Heads were isolated and brains were carefully dissected in ice-cold 1X phosphate-buffered saline (PBS) and moved into 4% paraformaldehyde (PFA) in 1X PBS. Brains were incubated in 4% PFA for 20 minutes and transferred to 0.5% Triton X-100 in 1X PBS (PBST) and washed four times (for at least 20 minutes per wash). Brains were then transferred to 5% normal goat serum (NGS) in PBST for one hour (blocking solution). For primary antibody staining we incubated brains overnight at room temperature on a shaker. Brains were then washed 4 times for 20 minutes per wash in PBST at room temperature. Brains were then transferred to the secondary antibody staining solution and incubated for three hours at room temperature on a shaker. Brains were then washed in PBST for four times for 20 minutes per each wash. Finally, brains were dehydrated with 25% glycerol then 50% glycerol in PBS for 30 minutes each. Finally, we mounted brains on poly-L-lysine covered slides (Sigma Aldrich) in Vectashield mounting medium (VectorLabs) and were kept in 4°C for at least one hour prior to imaging.

For paraFLPtag experiments we used polyclonal rabbit anti-RFP (abcam, ab62341; 1:5000) with polyclonal chicken anti-GFP (abcam, ab13970; 1:10000) in the primary antibody staining solution. For secondary antibodies we used goat anti-rabbit Alexa Fluor 633 (Invitrogen, A21070; 1:1000) and anti-chicken Alexa Fluor 488 (Invitrogen, A11039; 1:1000). For paramCherry, MB043C>UAS-myrGFP experiments we used rabbit anti-mCherry (abcam, ab167453 at 1:2,500); chicken anti-GFP (abcam, ab13970; 1:10,000). For secondary antibodies we used anti-rabbit AlexaFluor 633 (Invitrogen, A21070; 1:1000), and anti-chicken (Invitrogen, A11039; 1:1000).

#### Biocytin and Neurobiotin Staining

Neurobiotin and biocytin hydrazide staining was performed by carefully removing fly heads from the recording platforms and dissecting fly brains in 1X PBS. Brains were then transferred to 4% PFA in 1X PBS for 35 minutes at room temperature and kept in an aluminium foil wrapped well plate for the remaining steps. Brains were then washed in 1X PBS for 15 minutes at least three times. For neurobiotin staining we incubated brains at 4°C overnight with 1:400 streptavidin conjugated AlexaFluor 633 in 0.1% PBT. For biocytin hydrazide staining we incubated brains at 4°C for roughly 18 hours in 1:400 streptavidin conjugated AlexaFluor 633 in 0.1% PBT. We then washed brains in 1X PBS for 15 minutes three times and gently dehydrated brains for 1 hour in 30% glycerol in 1X PBS. Brains were mounted in Vectashield mounting medium and kept in 4°C for at least 1 hour prior to imaging.

#### Microscopy

Images were taken on a Zeiss LSM980 with Airyscan2 microscope (ZEN blue 3.3) with a Plan-Apochromat 63× (1.40 NA) oil immersion objective.

#### Biocytin Hydrazide

Imaging for α1 PAM DAN filled biocytin hydrazide images were taken using a Leica TCS SP5 X confocal microscope with an HCX IRAPO L x25 water immersion objective (0.95 NA). Images were taken at a resolution of 1024×512. Laser power was kept at 2% for Alexa-Fluor 633 channel and 1% for the GFP 488 channel.

#### Biocytin Reconstructions

Three-dimensional neuronal reconstructions were generated from the tdTomato channel for paraFLP tag experiments or AlexaFluor-633 channel for biocytin hydrazide experiments.

Image stacks were reconstructed using the Simple Neurite Tracer (SNT) Fiji plugin. Neuronal processes were traced in three dimensions using the interactive NBA* (New Bidirectional A*) path-search algorithm. For visualization, three-dimensional skeletons were aligned to the XY orientation corresponding to the acquired image stacks and exported from SNT as a Scalable Vector Graphic (SVG) file.

#### SIZ Reconstructions

As the MB043C-GAL4 labels multiple α1 DANs, we reconstructed the primary neurite branch containing all of the fasciculated processes. Para-GFP was mapped manually onto the tdTomato-derived neuronal skeleton. Because spike initiation zones (SIZ) with enriched Para-GFP expression was often only modestly elevated relative to baseline Para-GFP (outside of the expected SIZ), we could not apply automated intensity-based segmentation using a single fluorescence threshold to reliably distinguish SIZs. The transition between non-SIZ regions to SIZ was also frequently gradual rather than sharply defined, making the precise proximal and distal boundaries difficult to identify using a single threshold. SIZ boundaries were therefore assigned by manual inspection of the GFP channel throughout the three-dimensional image stack.

### Electrophysiology

*Fly tethering* was performed by briefly ice-anaesthetizing and tethering flies using UV-cured glue to a custom printed platform^70^. Using a small needle, we created a window by making cuts into the cuticle and carefully removing the cuticle, trachea, and fat bodies using forceps. To access the soma for patching we used a slightly broken pulled pipette (broken on a Kimwipe after pulled to 20-30 MΩ) to break through the glial sheath with mechanical force (positive pressure out of the broken pipette filled with extracellular saline). Pressure was monitored using a Manometer (RS Pro 273-9927).

Patch-clamp pipettes were pulled to a resistance of 13-15 MΩ for PAM recordings and 9-11 MΩ for MBON recordings using a DMZ Universal Electrode Puller (Zeitz). We recorded the spontaneous membrane potential for at least 5 minutes (Vm) in current clamp mode (I_clamp_ = 0) (Multiclamp 700B, Molecular Devices) at a sampling rate of 10 kHz (Digidata 1440a, Molecular Devices) and saved all recordings (pClamp software suite, Molecular Devices). Voltage recordings were corrected for a 13 mV junction potential.

For α1 PAM recordings, cells were given current injections ranging from steps of -10 pA to 8 pA in increments of +2 pA per step. For dual α1 PAM recordings cells this range was extended to – 20 pA to + 20 pA with increments of +5 pA. For α1 MBON recordings cells were given current injection steps from -30 to +30 pA in increments of +10 pA. For α1 DAN-MBON recordings, cells were recorded from -35 to +35 pA in increments of +5 pA.

*Biocytin* filled neurons were held for at least 1 hour when using neurobiotin tracer. For filling with biocytin hydrazide PAM DANs were filled for at least 4 hours or at least 1.5 hours if filling MBONs. A small holding current (∼5 pA for PAM and ∼10 pA for MBON) was applied once all recordings were obtained to increase biotin loading into neurons.

#### Bath Perfusion

For experiments with tetrodotoxin perfusion, we used a peristaltic pump (Watson-Marlow) to deliver external saline at a rate of 0.029 mL/sec, coupled with a custom vacuum-powered outlet. For drug perfusion we delivered at least 5 bath volumes of tetrodotoxin-laced extracellular saline to assure the maximum concentration was reached.

#### Extracellular saline

contained 103 mM NaCl, 3 mM KCl, 5 mM TES (N-Tris(hydroxymethyl) methyl-2-aminoethanesulfonic acid), 10 mM trehalose, 10 mM glucose, 2 mM sucrose, 26 mM NaHCO_3_, 1 mM NaH_2_PO_4_, 1.5 mM CaCl_2_, 4 mM MgCl_2_ (pH was equilibrated to 7.3 with 95% O_2_/5% CO_2_ and osmolarity was 275 mOsm). For sugar-free saline trehalose, glucose, and sucrose was replaced with equimolar ribose.

#### Internal saline

contained 140 mM potassium-aspartate, 1 mM KCl, 10 mM HEPES, 1 mM EGTA, 0.5 mM Na_3_GTP, 4 mM MgATP, potassium hydroxide (titrated to pH 7.3, 265 mOsm). Some pipettes also contained 13 mM biocytin hydrazide or 2% neurobiotin tracer with 0.2 mM Alexa Fluor™ 594 Hydrazide (Sigma A10438)^33^.

### Two-photon holography

#### Setup

To record SIZ1 activity while simultaneously activating SIZ2 and/or SIZ3, we used a holographic all-optical approach inspired from previous mice work^71–74^. The setup consisted of a Bergamo II two-photon microscope (Thorlabs) equipped with two femtosecond-pulsed lasers, enabling simultaneous two-photon stimulation and volumetric imaging of α1-DANs co-expressing GCaMP7f and ChrimsonR (*α1-DAN>ChrimsonR; GCaMP7f*). GCaMP7f signal was imaged at 920 nm using a Ti:Sapphire laser (MaiTai DeepSee, Newport Spectra-Physics), while ChrimsonR was stimulated at 1035 nm using an AeroPulse FS50 laser (Hamamatsu Photonics). Along the stimulation path, a 512×512-pixel spatial light modulator (Meadowlark Optics) and two galvanometric mirrors enabled three-dimensional targeting of SIZ2 and/or SIZ3 with diffraction-limited, spiral-shaped stimulation patterns. Along the imaging path, a remote focus device (Thorlabs) enabled simultaneous volumetric imaging of SIZ1 (13 planes spaced 5.5 µm) without physically moving the objective (x20 NA 1.0 XLUMPFLN, Olympus). To prevent stimulation artifacts, laser pulses were synchronized to the 8.3 kHz resonant scanner using ScanImage, restricting them to the scanner fly-back period.

#### Experimental procedure

Flies of the genotype α1-DAN>ChrimsonR; GCaMP7f were reared on food supplemented with all-trans retinal (400 μM final concentration, Cayman 116-31-4) and kept in darkness before testing. Brains from 6-8-day old flies were dissected in carbonated extracellular saline (ECS)^75^ and transferred to a poly-L-lysine-coated coverslip placed in an imaging chamber (ALAMS-518SWPW). Carbonated ECS was continuously perfused over the brain throughout the imaging session (Masterflex 78018-40). SIZ1 activity was imaged at 1.17 Hz while four stimulation conditions (2 s of 5 ms pulses at 100Hz; average power 1.6 mW/spiral) were delivered in randomized order: (1) SIZ2+3, (2) SIZ2 only, (3) SIZ3 only, and (4) no SIZ stimulation with stimulation spirals positioned away from the targeted SIZs. To avoid light aberrations associated with two-point holograms^76,77^, the stimulation pattern consisted of three spirals (each 5 µm in diameter, with 20 revolutions, 5 ms in duration). In the SIZ2+3 condition, two spirals were positioned one on SIZ 2 and one on SIZ 3. The third area was placed outside of the α1 DAN neurites. In the SIZ 2 or SIZ 3 only conditions, one spiral was positioned at the corresponding SIZ, while the remaining two spirals were positioned outside the α1 DAN neurites. Finally, in the control condition, all three spirals were positioned outside of the α1 DAN neurites. All imaging and stimulation parameters were controlled using ScanImage.

Changes in calcium fluorescence were calculated as ΔF/F = (F - F0)/F_0_, where F_0_ was the mean fluorescence during the 2 s preceding stimulation onset. Responses were quantified as the area under the curve during the stimulation period. Each session typically included two stimulations per condition. For ΔF/F_0_ analysis, stimulations within each condition were averaged across sessions for each brain. Data analysis was performed using Python and GraphPad Prism.

## Data Analysis

Electrophysiology recordings were analysed using a custom MATLAB script (version R2022a; Mathworks). For paired recordings two channels were analysed with identical procedures. Before analysis each voltage trace was band-stop filtered with a second-order Butterworth IIR filter (49-51 Hz, zero phase via filtfilt) to remove electrical noise. Subthreshold membrane potential was estimated by median filtering the raw trace (1 s window).

Spike detection was performed on a low-pass filtered signal (100 Hz). Candidate spikes were identified from the difference between low-pass filtered and median filtered traces to remove any slow drift. Then we computed the first derivative from this signal and spikes were detected as the peaks using a user-defined minimum peak prominence threshold. This threshold was set manually after visual inspection of the detection trace and the threshold criterion was recorded for each trace. Spike times were then refined for PAMs because slower spike kinetics of large amplitude spikes prevented accurate spike amplitude measurements. We therefore identified the local voltage maximum between the initial spike detection index and the afterhyperpolarization index following each spike. Duplicate indices were removed. Spike amplitude was defined as the difference between Vm at the spike time and the median-filtered baseline at that time. Inter-spike intervals were computed as the successive differences in spike times.

Bursts were defined as a group of at least three spikes separated by inter-spike-intervals below a threshold set for each recording by visual inspection. Thresholds were also recorded for each data file. Each burst was characterized by its onset (first spike), offset (last spike), duration (offset minus onset), and spike count, from which mean spikes per burst, burst duration, and burst rate were derived. IPSPs were detected from the difference between the median filtered baseline and the recorded trace (set to zero where negative so hyperpolarizing events appeared as positive deflections). Events were identified with MATLAB ‘findpeaks’ using a user-defined minimum peak prominence threshold after visual inspection. IPSP amplitude and inter-event intervals were extracted from the detected peaks. User defined thresholds for IPSPs were also recorded.

### Oscillatory Power Analysis

Spectral power was quantified from spontaneous activity recordings using ‘pwelch’ or Welch’s method. For each recording channels were band-stop filtered to remove electrical noise (49-51 Hz) and the first and last 2-3 seconds were removed to exclude termination artefacts from filtering. Traces were then down sampled to a working sampling rate matched to the band of interest (constrained to at least 4x the upper band to avoid aliasing) and linearly detrended. Power spectral density (PSD) was estimated with a Hamming-windowed Welch periodogram using 50% overlapping segments of 10 sec for 10 Hz and 60 sec for 1 Hz, with FFT length set to at least 8192 points to yield a fine frequency grid. Band power was computed as the summed PSD within target frequency range (±0.5 for 1 Hz or ± 2 for 10 Hz). For each recording and channel, band power was log_10_-transformed.

### Coherence Analysis

We computed the magnitude-squared coherence between two simultaneously recorded neurons across the full trace for each recording. Coherence was estimated using Welch’s averaged periodogram (MATLAB, mscohere) with a 10 second Hamming window, and 50% overlap and an FFT length set to the next power of two at or above the window length, with no additional zero-padding. The cross-power spectral density was estimated over the same segments (MATLAB, cpsd) and the phase of the cross-spectrum was retained as an estimate of the phase relationship between the two traces. From each coherence spectrum we extracted the mean and peak coherence. To determine the z-scored coherence we created 500 iterations of a circularly shifted surrogate (shifted at least 1 s), and coherence was recomputed against the unshifted partner. Z-scored coherence was then computed per frequency bin as the difference between the observed coherence and the mean of the surrogate distribution, divided by its standard deviation.

### Coincidence Analysis

Spike and IPSP coincidence between two simultaneously recorded cells were calculated for all events across the whole recording. Each event in a reference cell was scored as coincident if the event in the partner cell fell within a tolerance window (1 to 100 ms in 1 ms increments) and the coincidence probability was defined as the fraction of reference-cell events with a coincident partner. We computed this across each cell pair (cell 1 relative to cell2 and vice versa). To compute the z-scored coincidence values we used a circularly shifted null distribution that was generated by creating an event train in one of the cell recordings that was circularly shifted by a random offset (>500 ms). The coincidence was then recomputed and repeated with 1000 shifts to build a null distribution that preserved the number and within-train temporal structure of events. Z-score was computed as the observed probability minus the null mean, divided by the null standard deviation.

Coincidence was considered above chance when z exceeded 2.

### Connectome Analysis

Contacts between NPF+ neurons and α1 DANs (PAM11) were identified from visual inspection of the skeletons and EM slices (Flywire) in Fig. 1g. The coordinates for the EM images are x = 158907, y = 32349, z = 1656 for Fig. 1g, top and x = 2997625, y = 31145, z = 2857 for Fig. 1g, bottom. For Supplementary Fig. 2g,h, we obtained postsynaptic sites onto α1 DANs (PAM11) from the public FAFB dataset (fafbseg.flywire.get_synapses). Neuronal skeletons were obtained using the fafbseg.flywire Python interface from FlyWire materialization 783. Synapses were retrieved with filtered=False and a minimum cleft score of >50. Unidentified presynaptic root IDs were excluded. Synaptic inputs were also restricted to neurons that had at least 5 synapses between the pre-synaptic neuron and the α1 DAN. The resulting postsynaptic coordinates and corresponding postsynaptic neuron IDs were exported for further analysis.

For Supplementary Fig. 2g, Visualization of local input-synapse density was mapped onto the reconstructed neuronal mesh surface as a Gaussian kernel density estimate over three-dimensional Euclidean distance (bandwidth = 15 μm), evaluated at each mesh vertex. For every vertex, density was calculated as the sum of Gaussian-weighted contributions from all input synapses within three bandwidths, without further normalization; values are therefore reported in arbitrary units and used for visual and qualitative comparison across the arbor and across neurons within a single colour scale, rather than as an absolute density measure. Therefore, the surface estimate reflects three-dimensional spatial proximity to synapses across the whole reconstructed mesh, including the soma. Meshes were rendered with input-synapse density mapped to colormap “hot” (scale bar in arbitrary density units). Figures were generated using Plotly (Python) and rendered to static images at a fixed camera position.

Each postsynaptic site was mapped to its nearest skeleton node in three-dimensional space using a k-d tree (scipy.spatial.cKDTree). The origin for soma-rooted distance measurements was defined independently for each neuron as the skeleton node nearest to its annotated soma coordinate. Thus, distance zero corresponds to the skeleton node nearest the annotated soma position rather than to the nearest point on the soma surface.

*Geodesic cable distance* from this soma-associated skeleton node to all other skeleton nodes was calculated as the shortest path through the weighted skeleton graph using Dijkstra’s algorithm. Each input synapse was subsequently assigned a geodesic distance on its corresponding skeleton node. Consequently, synapse position on the plot represents the distance travelled along the reconstructed neuronal arbor from the soma-associated node and not the Euclidean distance between the soma and synapse.

For visualization, soma-rooted geodesic synapse distances were converted from nanometres to micrometres and plotted without normalization between neurons. Each PAM-α1 neuron was displayed as an individual row, and each retained postsynaptic site was represented by a vertical blue tick positioned at its measured geodesic cable distance from the soma-associated skeleton node. No kernel-density estimation, binning, interpolation, or spatial smoothing was applied to the synapse positions displayed in this panel; each tick therefore represents an individual postsynaptic site at its calculated absolute cable distance. Absolute rather than normalized distances were used so that differences in the physical extent and synapse distribution of individual neuronal arbors were retained. Three reference positions at 72.01, 154.2, and 160.0 µm from the soma-associated skeleton node were overlaid as vertical dotted lines for anatomical comparison.

### Statistical Analyses

Statistical analyses were performed in GraphPad Prism (10.6.1), MATLAB (R2022a), or Python (3.12.0). All behavioral, electrophysiological, calcium imaging, and confocal data were tested for normality using the Shapiro-Wilk test. Datasets not normally distributed were analysed using non-parametric equivalents. “Datasets were also checked for homogeneity of variance using the Brown–Forsythe test, and Welch’s correction was applied where variances differed. Finally, all tests were two-sided unless stated otherwise and significance was set at α = 0.05.

#### Repeated measures and multiple comparisons

Sphericity was not assumed for repeated-measures ANOVA and the Geisser-Greenhouse correction was applied throughout.

Repeated-measures datasets that did not pass the Shapiro–Wilk test were analysed with the Friedman test instead. All p values reported for multiple comparisons were corrected using the tests specified in the legends. Spearman correlation and linear regression are reported uncorrected.

*Correlations and regressions* were assessed using Spearman rank correlation for associations between physiological measures and reported as r_s_ and p values. Current-voltage relationships were fitted with a simple linear regression and reported as slope, R^2^, and p values.

*Coincidence based measures* were calculated as z-scores against circularly shifted surrogate distributions described above and coincidence was considered above chance when z>2.

*Sample size* No statistical method was used to predetermine sample size. Sample sizes were selected to be comparable to prior studies for in vivo patch clamp^26^ and behavior^7^.

#### Data exclusions

For whole cell patch-clamp recordings the recording quality was assessed by the seal quality, resting membrane potential, and recording stability. Recordings were performed such that spontaneous activity was first recorded for a few minutes then current injections were made to generate the IV plots. Some missing n for the IV plots are due to one or more cell being lost following the spontaneous activity recording. For power analyses the first and last 2-3 seconds of each trace were removed to exclude edge artifacts from the filter. For connectome analyses, synapses were retained at a minimum cleft score of 50 and inputs were restricted to presynaptic partners having ≥5 synapses onto the α1 DAN.

#### Replication

All electrophysiology, calcium imaging, confocal imaging, and behavioral experiments were performed across multiple days and experimental cohorts.

#### Randomization and blinding

Randomization was performed for holographic optogenetic stimulation experiments as the four stimulation conditions were delivered in a randomized order within each session. For behavioral tests genotypes were tested in a random order and CS+ odor identity was always counterbalanced across groups. For electrophysiology experiments groups were interleaved for starvation conditions and optogenetic stimulation protocols. Blinding was performed for manually set parameters of electrophysiology analyses, specifically burst analyses, spike and IPSP detection. Blinding was also performed for biocytin reconstructions. Experimenters were typically not blinded to conditions during the experimental procedures.

### Single-cell transcriptomic analysis

ScSeq data from the *Drosophila melanogaster* midbrain was downloaded (GEO: GSE207799^78^) and analyzed using Seurat in R. Dopaminergic neurons were isolated as cells co-expressing Vmat and DAT (normalised expression >4 and >3, respectively). These cells were re-normalised, scaled and re-clustered (50 PCAs, resolution = 4). *NPFR* expression was visualised on the UMAP embedding. α1-PAM DANs were identified by its expression of *Cht6* and *up*, which are markers of alpha1-PAMs according to bulk-RNA sequencing data of FACS samples in Aso et al.^38^. See supplemental material for code.

### Computational Model

We constructed a multi-compartmental conductance-based model of an α1 dopaminergic neuron in Brian 2^79^. Each compartment obeyed the cable equation as handled by Brian 2’s *SpatialNeuron*. Equations were integrated using the stochastic Heun method with a fixed time step of 1 ms. The somatic membrane potential was recorded throughout, and somatic spikes were detected from the resulting trace using a peak-finding algorithm (*scipy.signal.find_peaks*; minimum height −35 mV, minimum inter-peak separation 100 ms, minimum prominence 5 mV).

#### Morphology

The morphology consisted of a spherical soma (3 µm diameter) connected to a primary trunk (70 µm length, 0.25 µm diameter), which branched into an axonal arbour and a dendrite. The axon comprised a proximal segment (70 µm), a distal segment (130 µm), and a non-excitable collateral branch (200 µm); the dendrite was a single process (200 µm). All processes were 0.15 µm in diameter and were discretised at 1 µm spatial resolution. The specific membrane capacitance was *C_m_* = 10 µF cm⁻² and the axial resistivity *R_i_* = 25 Ω cm.

#### Membrane dynamics

The transmembrane current density was given by the sum of leak (*I_L_*), voltage-gated sodium (*I_Na_*), delayed-rectifier potassium (*I_K_*), voltage-gated calcium (*I_Ca_*), calcium-activated potassium (*I_SK_*), and calcium-activated non-selective cation (*I_CAN_*) currents:

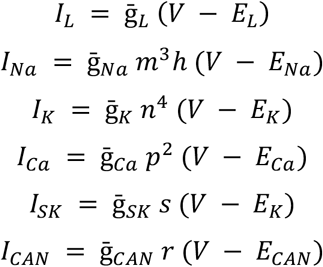

with total transmembrane current

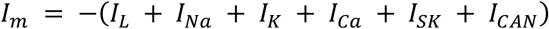

*Gating.* The voltage-gated activation and inactivation variables (*m*, *h*, *n*, *p*) followed first-order kinetics, *dx*/*dt* = −(*x*_∞_ − *x*)/*τ_x_*. Steady-state activations took the form

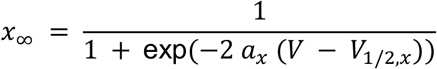

The calcium-gated activation variables (*s*, *r*) similarly followed first-order kinetics, *dy*/*dt* = −(*y*_∞_ − *y*)/*τ_y_*. Steady-state activations took the form:

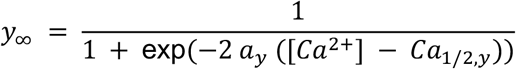

*Calcium dynamics.* Intracellular calcium was tracked as a single pool,

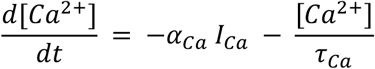

with influx scaled by *α_Ca_* and decay scaled by time constant *τ_Ca_*.

#### Spatial distribution of conductances

Conductances were distributed to create three spatially restricted spike initiation zones (SIZs), positioned to match the para-enriched regions identified anatomically. The soma and trunk carried sodium and potassium conductances at 5% of their maximal density, and the remaining non-SIZ axonal and dendritic membrane carried them at 4%; the collateral branch was passive (no Na or K). Against this low-excitability background, three SIZs were endowed with elevated conductances: a proximal-axonal (branch) SIZ (0 µm to 20 µm from the branch point) with sodium and CAN at 150%, and full potassium and SK density; a distal-axonal SIZ (90 µm to 130 µm from the branch point) with full sodium and potassium and no SK; and a dendritic SIZ (85 µm to 175 µm from the branch point) with sodium and potassium at 50% and no SK. SK conductance was additionally placed on the membrane proximal to the axonal and dendritic SIZs. Calcium conductance was uniform across the neuron. The branch SIZ was therefore the only zone equipped with both of the calcium-activated SK and CAN currents that support burst and plateau dynamics.

#### Stimulation and noise

Synaptic-like drive was delivered as point currents to 5 µm segments centred on the dendritic and axonal SIZs. Inputs were Poisson trains (0.5 Hz at each site) of 50 ms, 5 pA current pulses, delivered over a 140 s window flanked by 500 ms of baseline. Inter-event intervals were drawn from an exponential distribution. Ornstein–Uhlenbeck noise (with σ = 1 pA, τ = 300 ms) was injected at the soma.

## Data Availability

Sample electrophysiology, calcium imaging data, confocal stacks, and behavioral data generated in this study will be deposited in DOI: **10.5281/zenodo.23197282.** Connectomics analyses used publicly available FlyWire FAFB data (https://flywire.ai).

## Code Availability

Custom code for biophysical modelling can be found at https://github.com/ianconehed/hh_multi_spike_model; for connectomics analysis https://github.com/anniepark0/Connectomics-Geodesic-Distance-Code; for single cell transcriptome feature plot generation https://github.com/anniepark0/Single-Cell-NPFR-Feature-Plot; and for electrophysiology analyses https://github.com/anniepark0/Electrophysiology-Analysis.

## Supplementary Figure Legends

**Supplementary Figure 1.**
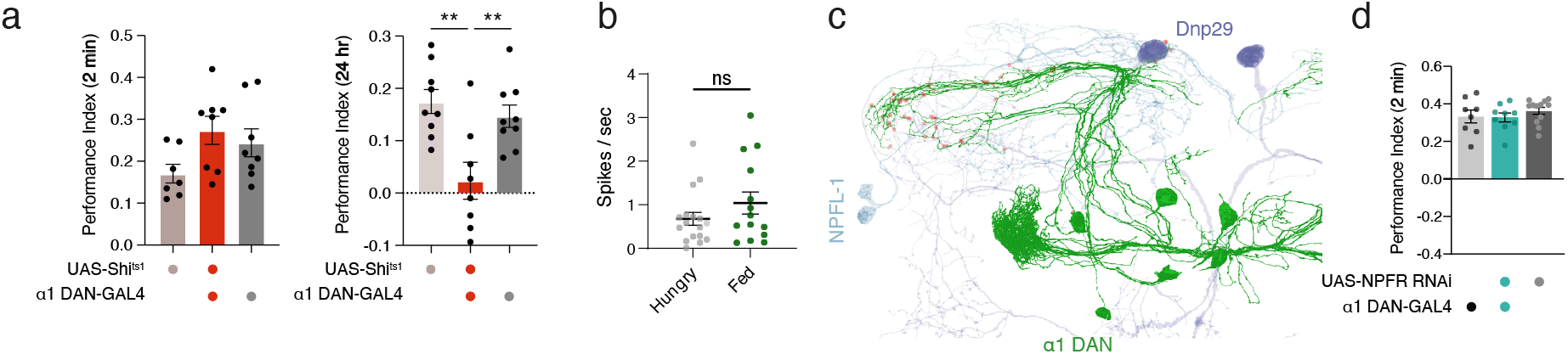
**a,** Inhibiting α1 DANs during training suppresses performance for long-term sucrose memory, but not short-term memory (STM n=7,8,8 and LTM n = 9,8,9 groups of flies (left to right), Tukey’s multiple comparisons test; UAS-Shi^ts1^ x α1 DAN-GAL4 (MB043C-GAL4) vs. α1 DAN-GAL4 (MB043C-GAL4)/+ p=0.0095, UAS-Shi^ts1^ x α1 DAN-GAL4 (MB043C-GAL4) vs. UAS-Shi^ts1^/+ p=0.0016). **b,** Spike frequency is not different between hungry and fed control flies (Mann-Whitney test; p=0.3771, n =17,14 neurons – one per fly). **c,** 3D Reconstructions of NPF expressing neurons (dark blue, Dnp29 and light blue, NPFL-1 or LNd) with α1 DANs (green) and sites of close proximity between them (red dots) (FAFB). **d,** NPFR knockdown does not alter short-term memory performance (Kruskal-Wallis with Dunn’s multiple comparisons test, n=8,9,12 groups of flies).

**Supplementary Figure 2.**
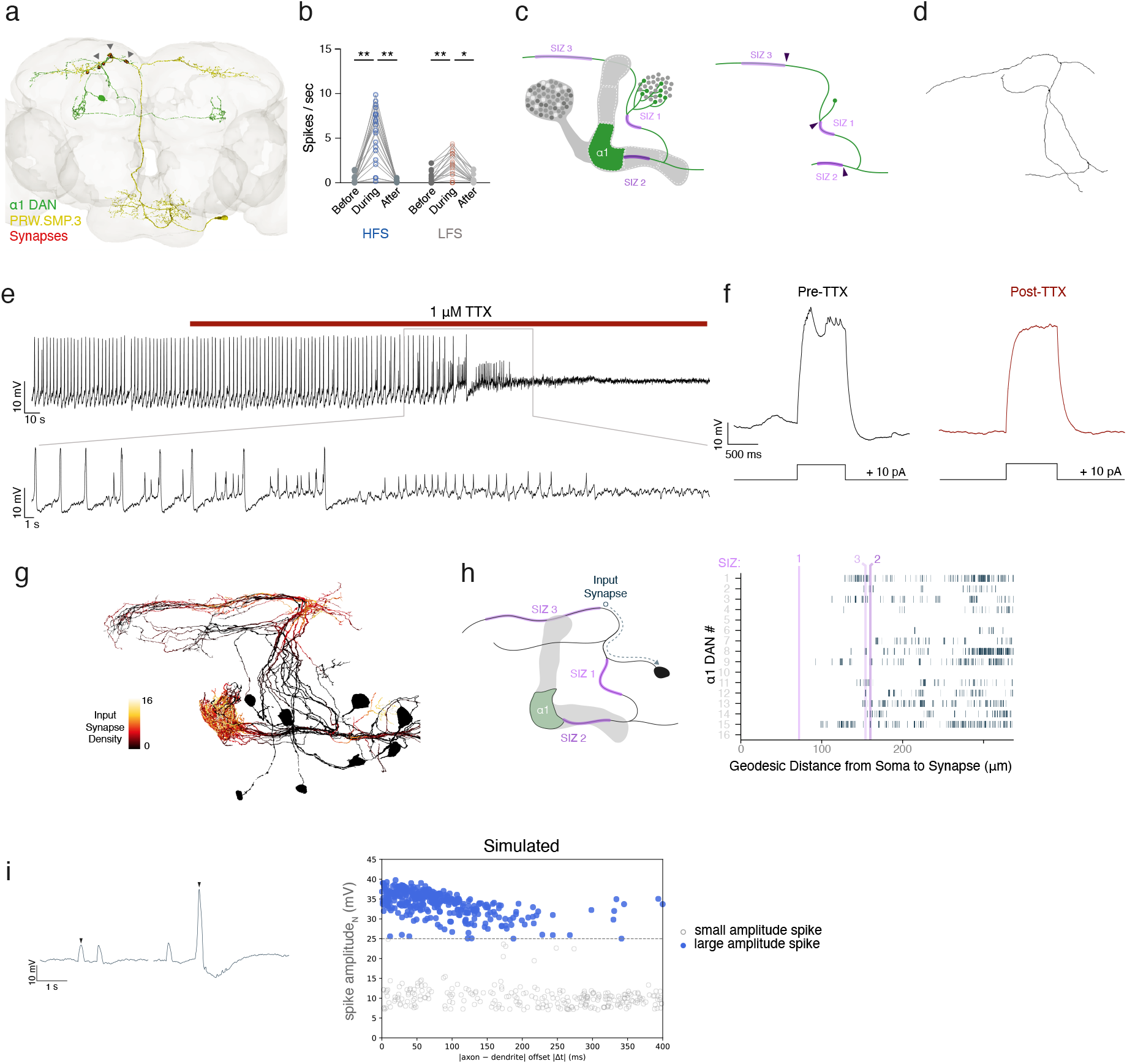
**a,** EM reconstructions of a secondary taste projection neuron (PRW.SMP.3, yellow) projecting onto the dendrites of a representative α1 DAN and the synapses between them (red, grey arrowhead) (FlyWire). **b,** Spikes of both large and small amplitudes elicited before, during, and after HFS or LFS of *Gr43a*-expressing primary taste neurons. Both HFS and LFS elicit an increase in spike frequency during optogenetic stimulation (n=20 neurons – 1 neuron per fly – with repeated measures; Friedman test with Dunn’s multiple comparisons, HFS Before vs. During p < 0.0001, During vs. After p < 0.0001; LFS Before vs. During p = 0.0043, During vs. After p = 0.0248). **c,d** Manual reconstructions were performed on FlpTag para and tdTomato labelled α1 DANs (Figure 2e). Reconstructions of the neurons were done on the main branches and distances to SIZ were calculated as the distance from the edge of the SIZ closest to the soma (black arrows). **e,** Representative trace of a large amplitude spiking cell perfused with 1 micromolar tetrodotoxin (TTX). Both large and small amplitude spikes subside with TTX perfusion. **f,** α1 DAN producing both spike amplitudes before tetrodotoxin (Pre–TTX) loses all spiking after TTX perfusion (Post-TTX). **g,** Input synapse density onto α1 DANs (Gaussian KDE, bandwidth=15 µm; FAFB). **h,** Left, geodesic distance (dotted line) from soma to an input synapse (circle). Right, geodesic distances per synapse across α1 DANs (rows). Magenta lines are SIZ distances from Fig. 2f. **i,** Left, Expanded segments of the simulated somatic voltage trace showing two characteristic events from the model: a small-amplitude spike (left) and a large-amplitude spike (right). Right, simulated somatic spike amplitude as a function of the interval between inputs delivered to SIZ 2 and SIZ 3. Large-amplitude spikes are more likely to arise when the two inputs arrive less than ∼150 ms apart; at longer intervals, the inputs tend to produce small-amplitude events.

**Supplementary Figure 3.**
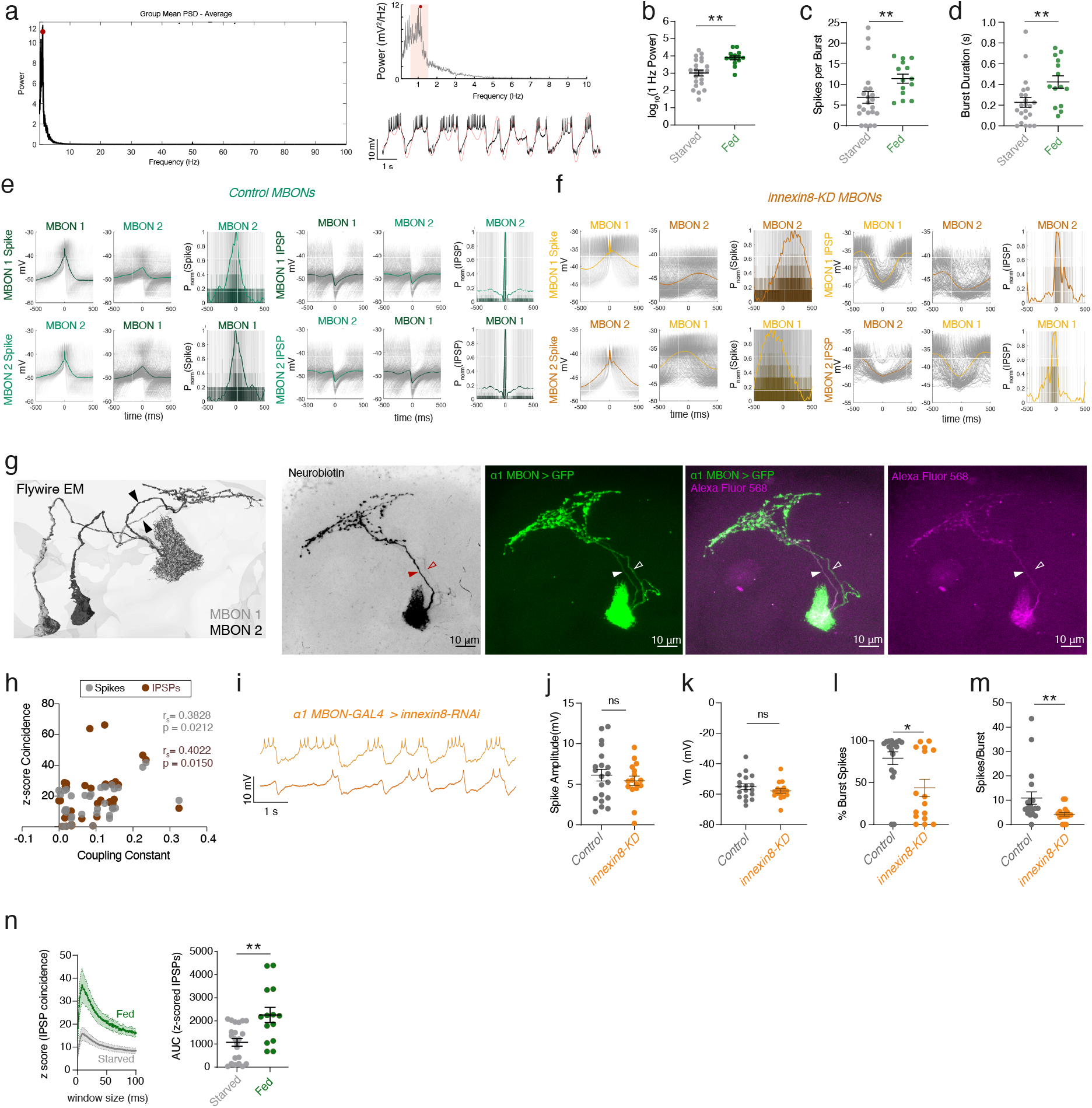
**a,** Left, Average power spectrum density plot for α1 MBONs. Top right, zoomed in power spectrum density plot from 0.1 to 10 Hz. Red dot indicates maximum value, and shaded red indicates 0.5-1.5 Hz range. Bottom right, example trace in black with superimposed filtered signal (filtered from 0.5 to 1.5 Hz). **b,** 1 Hz Power is greater in α1 MBONs of fed flies compared to starved flies (Mann-Whitney test; p=0.0008, n=14,22 neurons – 2 per fly). **c,** Number of spikes per burst increases in α1 MBONs from fed vs. starved flies (Mann-Whitney test; p=0.0049, n=14,22 neurons – 2 per fly). **d,** Burst duration increases in α1 MBONs of fed vs. starved flies (Mann-Whitney test; p=0.0084, n=14,22 neurons – 2 per fly). **e,** Left, spike-triggered averages of control MBONs from Fig. 3c. Right, IPSP-triggered averages between the same α1 MBONs. **f,** Left, representative spike-triggered averages for an α1 MBON pair with *innexin8*-KD. Right, IPSP triggered averages for the same pair. **g,** Left, EM reconstructions of ipsilateral α1 MBONs. Neurites of ipsilateral α1 MBONs overlap except for the neurite area labelled with black arrowheads. Left middle, image of α1 MBON directly filled with neurobiotin (filled arrowhead) and the ipsilateral α1 MBON was also labelled (hollow arrowhead). Right, only the patched cell contains AlexaFluor 568. **h,** Coupling constant correlates with z-scored spike (grey dots, Spearman correlation; r=0.3828, p=0.0212) and IPSP (brown, Spearman correlation; r=0.4022, p=0.015) coincidence (n=36 individual neurons – 2 per fly). **i,** Representative voltage trace of α1 MBON recording with *innexin8-*KD (MB310C>innexin8-RNAi). **j,** Spike amplitude (Welch’s test; p=0.5258, n=20,16 neurons – 2 per fly), and **k,** resting membrane potential (Welch’s test; p=0.2554, n=18,16 neurons – 2 per fly) are not different between control flies and *innexin8*-KD. **/:** Proportion of burst spikes is reduced in *innexin8*-KD (Mann-Whitney test; n=18,16 neurons, p=0.0147). **C:** Number of spikes per burst is reduced in *innexin8-*KD (n=18,16 neurons, Mann-Whitney test, p=0.0007). **2:** Fed flies exhibit greater coincident IPSPs than starved flies. Left, average z-scored IPSP coincidence curve across flies (mean ± s.e.m.) for fed (green, n=14 neurons) vs. starved (grey, n=22 neurons) flies. Right, AUC for individual z-scored IPSP coincidence curves (Mann-Whitney test; p=0.0027, n=14,22 neurons – 2 per fly).

**Supplementary Figure 4.**
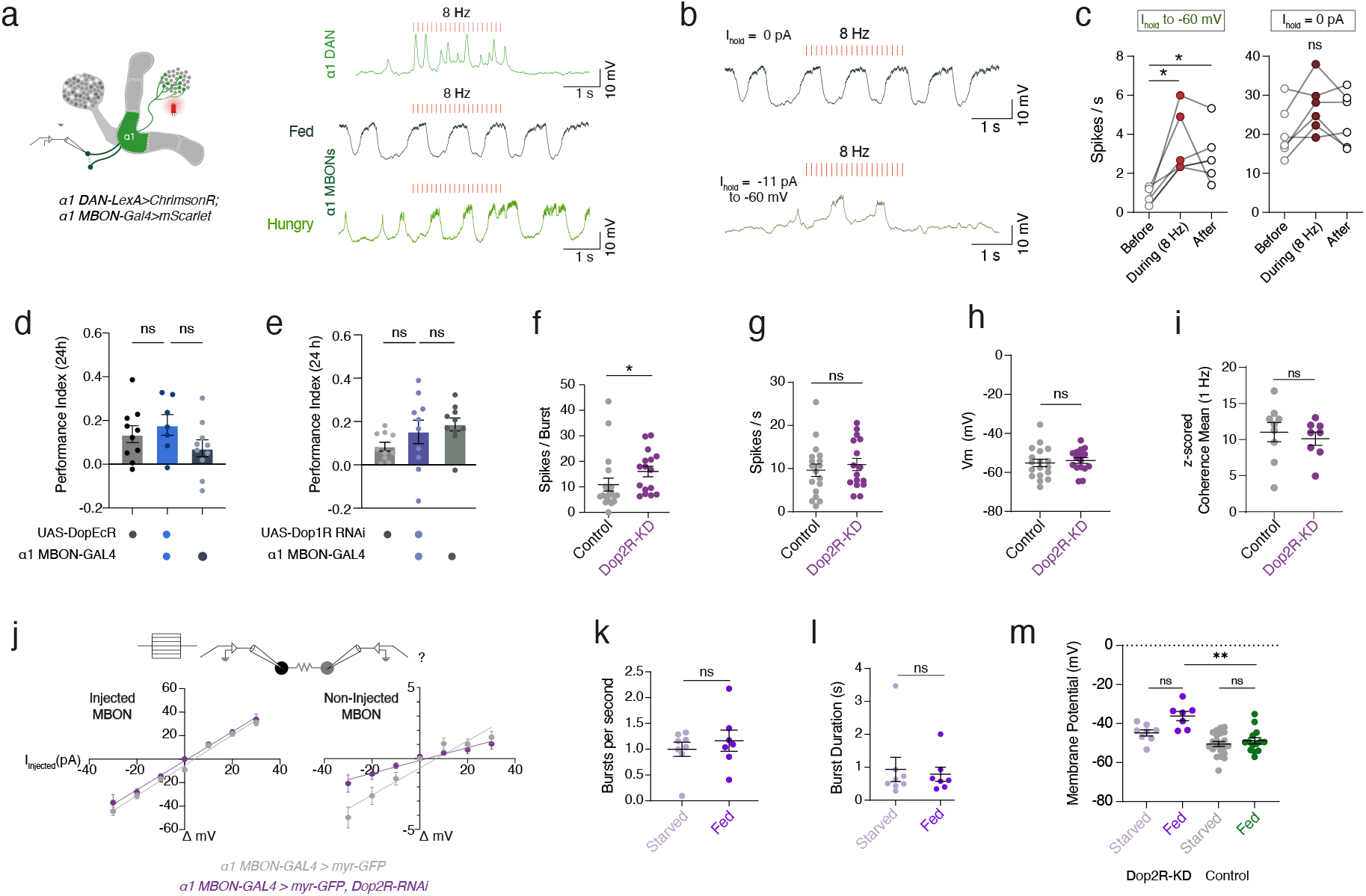
**a,** Traces of a representative α1 DAN receiving 8 Hz optogenetic stimulation (top) aligned with α1 MBON recordings (middle and bottom) to show minimal α1 DAN stimulation within physiologically relevant levels able to exhibit large amplitude spikes. **b,** Representative trace of α1 MBONs during α1 DAN stimulation with no current injected (top) or hyperpolarizing current injected to reduce resting membrane potential to -60 mV (bottom). **c,** Hyperpolarized α1 MBONs exhibited a change in spike rate (left; Friedman test; Before vs. During p=0.0187, Before vs. After p=0.0187, n=6), whereas those without current injection did not (right; Friedman test; Before vs. During p=0.4964, Before vs. After p=0.2978, n=6 neurons, one per fly). **d,** *29FHI$* knockdown did not change LTM performance compared to controls (n=10,7,10 groups of flies, Tukey’s multiple comparisons, p>0.05). **e,** *29F1$* knockdown did not affect LTM compared to controls (n=10 groups of flies, Dunn’s multiple comparisons test, p>0.05). **f,** Spikes-per-burst (Mann-Whitney test; p=0.0167, n=18,16 neurons – 2 per fly) increased in the *29FG$*-KD compared to controls. **g,** However, all spikes did not increase compared to controls, indicating spikes are reorganized into a bursting pattern (Mann-Whitney test; p=0.5281, n=18,16 neurons – 2 per fly). **h,** Membrane potential is not different between *29FG$*-KD and control flies (Mann-Whitney test; p=0.4426, n=18,16 neurons – 2 per fly). **i,** Z-scored 1 Hz mean coherence is not different between *29FG$\**KD and controls (Mann-Whitney test; p=0.6334, n=9,8 pairs of neurons, one per fly). **j,** I-V plot for control (grey) and *29FG$*-KD (purple) for injected MBON (left) and the resultant change in voltage in the non-injected MBON (right). Error bars are s.e.m. (Control n=7 pairs of neurons, Dop2R n=10 pairs of neurons). *29FG$*-KD suppresses coupling between pairs of α1 MBONs compared to controls. **k,** Bursts-per-second (Mann-Whitney test; p=0.8665, n=8,7 neurons – one per fly) and **l,** burst duration are not different between fed or starved *29FG$*-KD flies (Mann-Whitney test; p=0.9551, n=8,7 neurons – one per fly). **m,** Membrane potential does not differ between starved vs. fed *29FG$*-KD or control flies. However, fed *29FG$*E2* flies are more depolarized than fed controls (Kruskal-Wallis Dunn’s multiple comparisons test; p = 0.0014, n = 8,7,22,14).

**Supplementary Figure 5.**
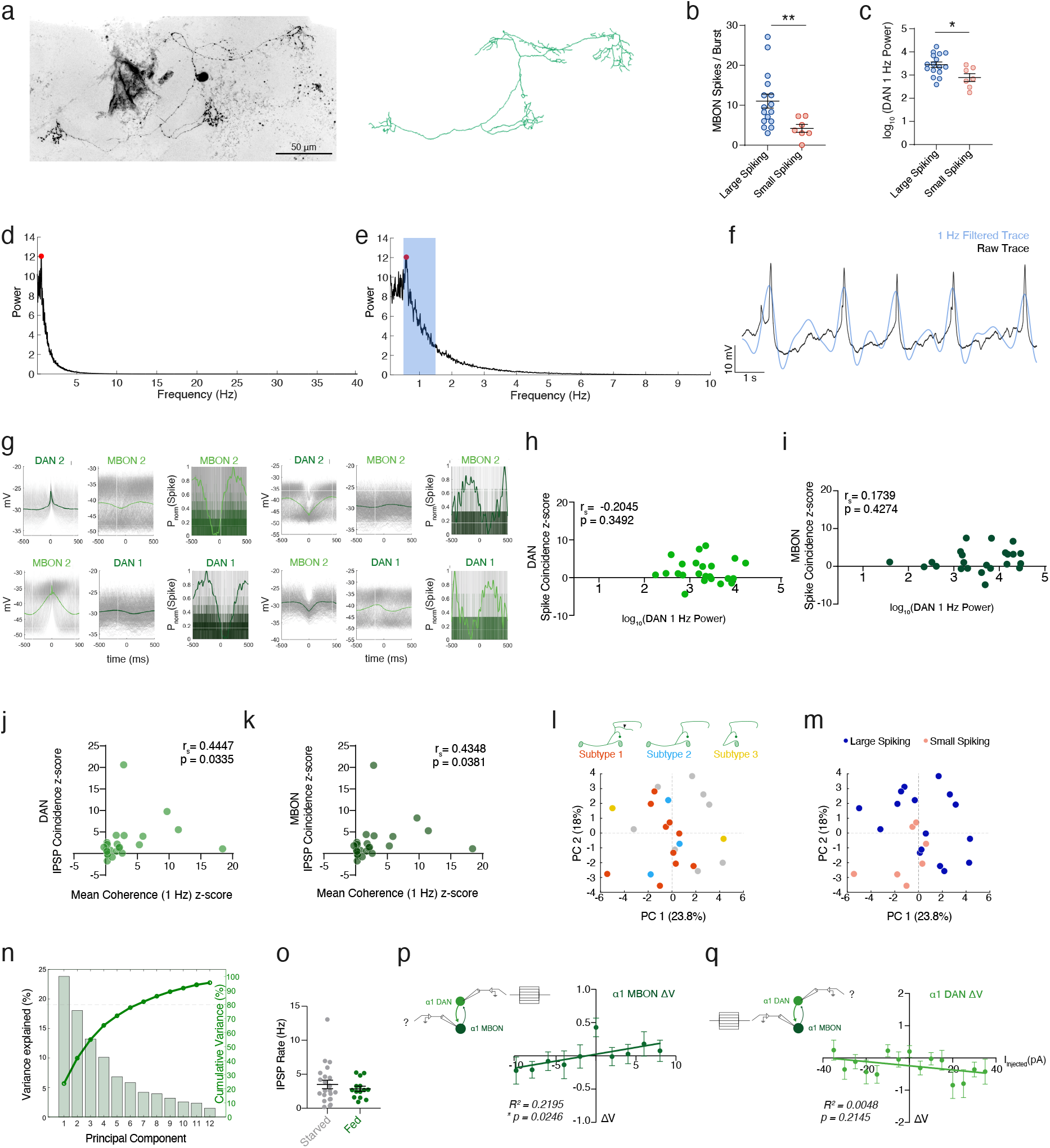
**a,** Left, representative biocytin filled α1 DAN and reconstruction (right). **b,** Number of spikes-per-burst in α1 MBONs is greater when α1 DAN exhibits large amplitude spikes (Mann-Whitney test; p=0.0075, n=16,7 neurons – one DAN-MBON pair per fly). **c,** DANs that can generate large amplitude spikes have greater power in the 1 Hz range (0.5 – 1.5 Hz) (Welch’s test; p=0.0187, n=16,7 neurons – one DAN-MBON pair per fly). **d,e,** Power spectrum density across α1 DAN recordings. **f,** Example raw electrophysiology trace (black) of an α1 DAN and an overlaid 1-Hz range bandpass filtered trace (blue). **g,** Left, spike-triggered average of DAN-MBON pair with anti-correlated spiking and IPSPs (right). **h,** 1 Hz power of individual DAN does not correlate with z-scored spike coincidence of DANs (Spearman correlation; n=23 DAN-MBON neuron pairs – one pair per fly, r=-0.2045, p=0.3492), or **i,** MBONs (Spearman correlation; n=23 DAN-MBON neuron pairs – one pair per fly, r=0.1739, p=0.4274). **j,** Z-scored 1 Hz mean coherence between α1 DAN-MBON pairs correlated significantly with Z-scored DAN IPSP coincidence (Spearman correlation; n=23 DAN-MBON neuron pairs – one pair per fly, r=0.4447, p=0.0335) and **k,** MBON IPSP coincidence (Spearman correlation; n=23 DAN-MBON neuron pairs – one pair per fly, r=0.4348, p=0.0381). **l,** PCA of all dual recorded pairs colored by anatomical subtype. Subtypes are as defined in Li et al. (2020). Subtype 1=α1-aad, Subtype 2=α1-dd, Subtype 3=α1-nc (n = 23 DAN-MBON neuron pairs – one pair per fly). **m,** The same PCA with individual pairs coloured by DAN spiking mode. **n,** Variance explained from top three principal components explains 23.8%, 18%, and 13.2% variance in the dataset. **o,** IPSP rate of α1 MBONs is not different between starved and fed flies (Mann-Whitney test; p=0.9362, n=22,14 neurons – 2 neurons per fly). **p,** Current injection into DANs evokes excitatory responses in MBONs (Linear regression; Slope=0.02044, R^2^=0.02195, p=0.0246, n=23 DAN-MBON neuron pairs – one pair per fly). **q,** Current injection into MBONs evokes a non-significant inhibitory response in DANs (Linear regression; Slope=-0.004881, R^2^=0.004811, p=0.2145, n=23 DAN-MBON neuron pairs – one pair per fly).

**Supplementary Figure 6.**
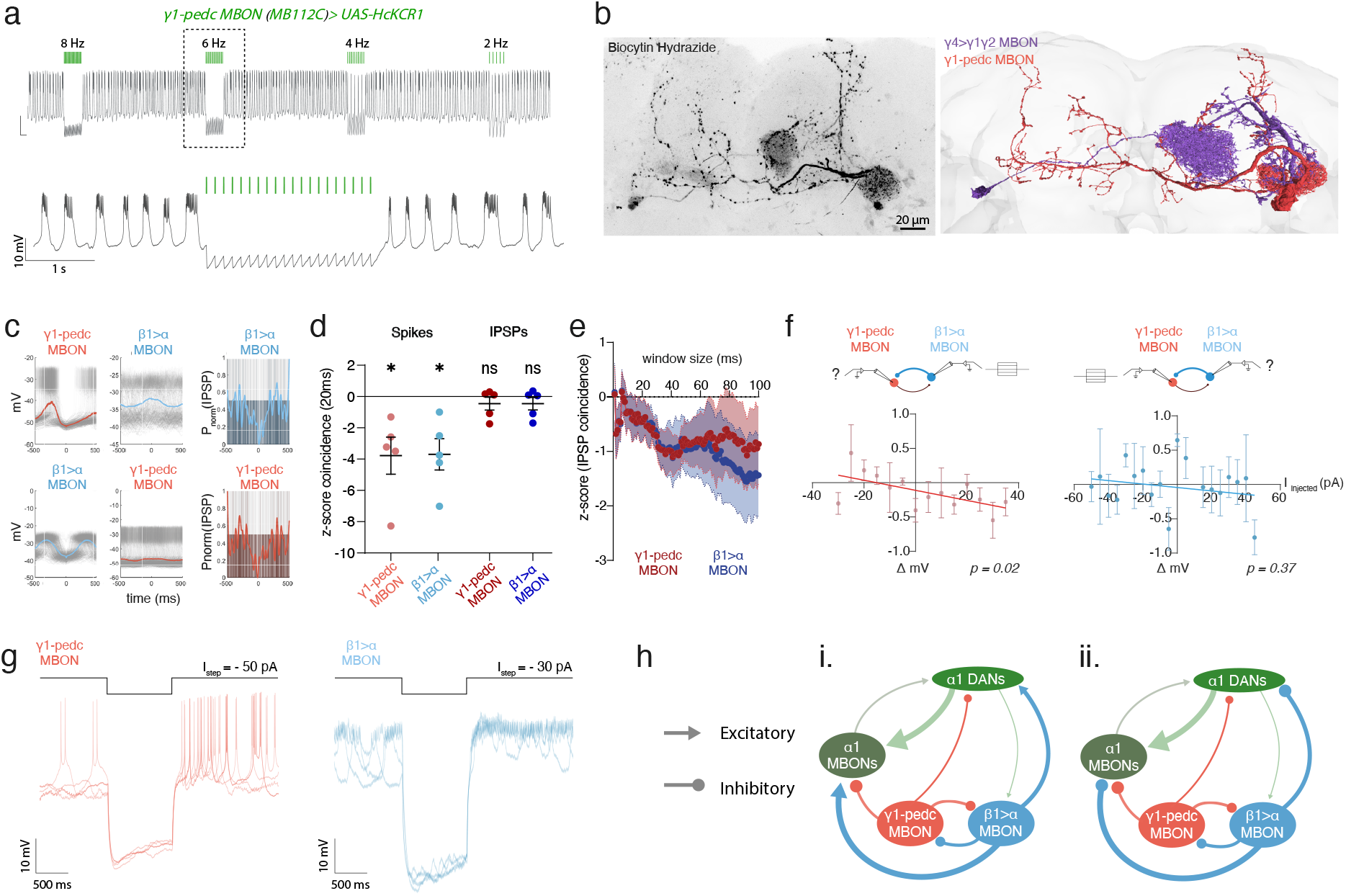
**a,** Representative trace of Y1-pedc MBON with optogenetic inhibition using *JIE4$* at different frequencies (5 ms pulse width, 2.5 second pulse train). **b,** Left, representative image of a biocytin labelled pair of Y1-pedc MBON and Y4>Y1Y2 MBON and their equivalent EM reconstructions (right). **c,** IPSP triggered average for a representative trace for a Y1-pedc MBON and β1>α MBON. **d,** Z-scored spike coincidence is significantly below average in Y1-pedc MBON and β1>α MBON, further supporting the idea they form an antiphase oscillator (One-sample t-test µ=0; Y1-pedc MBON p= 0.0335, p=0.0209 β1>α MBON, n=5 Y1-pedc and β1>α MBON pairs, one per brain). However, z-scored IPSP coincidence is not different from 0 (One-sample t-test µ=0; Y1-pedc MBON p=0.3293, p=0.3135 β1>α MBON, n=5 Y1-pedc and β1>α MBON pairs, one per brain). **e,** Z-scored IPSP triggered average between Y1-pedc MBON and β1>α MBON plotted across different window sizes. Shaded error is SEM (n = 5 Y1-pedc and β1>α MBON pairs, one per brain). **f,** I-V plots for the non-injected cell between dual Y1-pedc MBON and β1>α MBON recordings. Left, current injection into β1>α MBON and resultant change in voltage of Y1-pedc MBON, reveals that β1>α MBON inhibits Y1-pedc MBON (Linear regression; Slope=-0.007359, R^2^= 0.06606, p=0.0183, n=5 β1>α MBONs, one per brain). Right, current injection into Y1-pedc MBON and resultant change in voltage of β1>α MBON reveals no significant coupling between pairs (Linear regression; Slope=-0.002509, R^2^= 0.01022, p=0.372, n=5 β1>α MBONs, one per brain). **g,** Overlaid example traces from one sample Y1-pedc MBON and β1>α MBON with hyperpolarizing current injections showing post-inhibitory rebound. **h,** Putative wiring diagrams of the functional circuit connectivity if (i) glutamatergic β1>α MBON is excitatory (e.g. NMDAR expression in α1 MBONs, but GluClα is expressed in Y1-pedc MBON) providing the depolarizing phase of bursting in α1 MBONs, or if (ii) glutamatergic β1>α MBON is inhibitory (e.g. GluClα is expressed in α1 MBONs) but the kinetics of inhibition are slower or faster than GABAergic inhibition which provides in-phase inhibition, deepening the hyperpolarization phase of the α1 MBONs. The other possibility from motif (ii) is that the magnitude of inhibition from β1>α MBONs is smaller and only has an effect in sharpening inter-spike-interval within bursts.

